# Modelling the structure of Short Gastrulation and generation of a toolkit for studying its function in *Drosophila*

**DOI:** 10.1101/2022.05.13.491850

**Authors:** Sophie L Frampton, Catherine Sutcliffe, Clair Baldock, Hilary L Ashe

## Abstract

A BMP gradient is essential for patterning the dorsal-ventral axis of invertebrate and vertebrate embryos. The extracellular BMP binding protein Short Gastrulation (Sog) in *Drosophila* plays a key role in BMP gradient formation. In this study, we combine genome editing, structural and developmental approaches to study Sog function in *Drosophila*. We generate a *sog* knockout fly stock, which allows simple reintegration of altered versions of the *sog* coding sequence. As proof-of-principle, we test the requirement for two cysteine residues that were previously identified as targets for palmitoylation, which has been proposed to enhance Sog secretion. However, we show that the SogC27,28S mutant is viable with only very mild phenotypes, indicating that these residues and their potential modification are not critical for Sog secretion *in vivo*. Additionally, we use experimental negative stain EM imaging and hydrodynamic data to validate the AlphaFold structure prediction for Sog. The model suggests a more compact shape than the vertebrate ortholog Chordin and conformational flexibility between the C-terminal von Willebrand C domains. We discuss how this altered compactness may contribute to mechanistic differences in Sog and Chordin function during BMP gradient formation.

**Summary statement:** The authors model the structure of Sog and establish a new *sog* knockout fly stock that they validate for the testing of specific *sog* mutations.

## Introduction

Bone Morphogenetic Proteins (BMPs) are a large class of highly conserved signalling molecules that belong to the TGF-beta superfamily. BMPs perform essential roles during animal development and adult tissue homeostasis, the significance of which is reflected in the variety of human diseases attributed to aberrant BMP activity (Bandyopadhyay et al., 2013; Wang et al., 2014). BMPs bind to their receptors resulting in the phosphorylation of a receptor regulated Smad, which then forms a complex with the common mediator Smad that accumulates in the nucleus to regulate target gene transcription (Schmierer & Hill, 2007). A key developmental role for BMPs is the patterning of the dorsal-ventral (DV) axis in early vertebrate and invertebrate embryos. BMP gradient formation is mediated by a conserved network of regulators, including two BMP binding proteins, Short Gastrulation (Sog)/Chordin and Twisted Gastrulation (Tsg), as well as a protease, Tolloid (Tld) (Madamanchi et al., 2021; Matsuda et al., 2016; Shilo et al., 2013).

The most potent BMP signalling molecule in the early *Drosophila* embryo is a heterodimer of the BMP ligands Decapentaplegic (Dpp) and Screw (Scw) (Shimmi *et al*., 2005), which have uniform expression in the dorsal ectoderm (St Johnston and Gelbart, 1987; Arora *et al*., 1994; Shimmi *et al*., 2005). *tsg* and *tld* are also expressed in the dorsal ectoderm, while *sog* is expressed ventro-laterally in the neuroectoderm (Francois et al., 1994; Marqués et al., 1997; Mason et al., 1994). During embryogenesis, reciprocal gradients of Dpp/Scw and Sog are established across the dorsal ectoderm (Ferguson and Anderson, 1992; Ashe and Levine, 1999; Srinivasan *et al*., 2002; Shimmi *et al*., 2005; Wang and Ferguson, 2005). A narrow stripe of peak BMP signalling occurs along the dorsal midline and is flanked by lower signalling levels (Dorfman and Shilo, 2001; Rushlow *et al*., 2001; Sutherland *et al*., 2003; Shimmi *et al*., 2005) thereby subdividing the dorsal ectoderm into amnioserosa and dorsal epidermis, respectively (Raftery & Sutherland, 2003).

A favoured model of BMP gradient formation requires the shuttling of BMP ligands dorsally in a multi-protein complex (Holley *et al*., 1996; Marqués *et al*., 1997; Eldar *et al*., 2002; Shimmi *et al*., 2005; Umulis *et al*., 2006, 2009; Sawala *et al*., 2012). The model proposes that secreted Dpp/Scw binds to the extracellular matrix protein collagen IV (Col IV), which acts as a scaffold to promote formation of a Dpp/Scw-Sog-Tsg complex (Sawala et al., 2012; Wang et al., 2008). In this inhibitory complex, the Dpp/Scw ligand is unable to interact with its receptors but can diffuse dorsally (Ross *et al*., 2001; Eldar *et al*., 2002; Shimmi *et al*., 2005; Sawala *et al*., 2012). Cleavage of Sog within this complex by Tld liberates Dpp/Scw, allowing the ligand to re-bind Col IV. In dorso-lateral regions, close to the *sog* expression domain, the Dpp/Scw-Sog-Tsg complex is reassembled, resulting in inhibition of signalling and diffusion of the complex towards the dorsal midline. At the dorsal midline and in the absence of Sog, however, Dpp/Scw is free to interact with receptors, resulting in a graded BMP signal across the dorsal ectoderm (Sawala et al., 2012; Wang et al., 2008; Winstanley et al., 2015).

Sog function is also important in *Drosophila* pupal wing vein patterning, including formation of the posterior crossvein (PCV), which depends on signalling by Dpp and Glass bottomed boat (Gbb) ligands (Serpe et al., 2005; Wharton et al., 1999; Yu et al., 1996). As in embryogenesis, Sog functions with a Tsg-like protein, Crossveinless (Cv), and a Tolloid-related (Tlr) metalloprotease to both locally inhibit BMP signalling and enhance it at a distance from the source in the pupal wing (Ralston & Blair, 2005; Serpe et al., 2005; Vilmos et al., 2005). In this model, Dpp/Gbb is transported from the longitudinal veins in a Dpp/Gbb-Sog-Cv complex to the presumptive PCV, where it is released from the inhibitory complex by Tlr-mediated Sog cleavage, enabling ligand-receptor interactions (Shimmi *et al*., 2005; Serpe *et al*., 2005).

Sog and its vertebrate ortholog Chordin each contain four cysteine rich von Willebrand type C (vWC) domains which mediate protein interactions. These domains are 60-80 residues in length and have been identified in approximately 500 extracellular matrix proteins (Garcia Abreu et al., 2002; O’Leary et al., 2004; Zhang et al., 2007). Sog/Chordin vWC1 is separated from vWC2/3/4 domains by a ‘stem’ region comprising four Chordin specific (CHRD) domains (Francois et al., 1994). Structures have been solved for the human Procollagen IIA and the zebrafish Crossveinless-2 vWC1 domains (O’Leary et al., 2004; Xu et al., 2017; Zhang et al., 2008), however there is currently no experimental structure for these domains in Sog, or the Sog/Chordin specific 4x CHRD ‘stem’ region.

Sog secretion is critical to its function, and a previous study has reported a potential role for palmitoylation during this process (Kang & Bier, 2010). Palmitoylation is a lipid modification that can influence protein interactions, membrane association, and trafficking between sub-cellular compartments (Bannan et al., 2008; Kang & Bier, 2010; Linder & Deschenes, 2003). The *Drosophila* palmitoyl-transferase Huntington-interacting protein 14 (dHIP14) was identified as an interacting partner of Sog, and Sog was shown to be palmitoylated in tissue culture cells (Giot et al., 2003; Kang & Bier, 2010). Mis-expression of dHIP14 in embryos and wings reduced BMP activity, similar to Sog overexpression phenotypes (Giot et al., 2003; Kang & Bier, 2010). In addition, mutation of cysteines 27 and 28 of Sog prevented the dHIP14-mediated increase in Sog secretion in tissue culture, suggesting that these two residues are the primary palmitoylation targets (Kang & Bier, 2010).

In this study, we generate a *sog* knockout ‘reintegration-ready’ fly stock that we use to test the effect of mutating the palmitoylation sites *in vivo*. Our data show that these residues are not critical for Sog function. In addition, we combine EM imaging and Alphafold prediction to model Sog structure, which reveals a curved compact shape. This Sog structure, along with the *sog* knockout fly stock that we describe, will facilitate a complete molecular dissection of this critical extracellular BMP regulator.

## Results

### Generation of a *sog* KO with CRISPR

To facilitate analysis of Sog *in vivo*, we used CRISPR genome editing to generate a *sog* knock-out line in which the translation start codon of the endogenous *sog* locus on the X chromosome was replaced with a PhiC31 recombination landing site (Baena-Lopez et al., 2013) (Fig. 1A). Specifically, CRISPR-Cas9 mediated homology directed repair (HDR) was used to delete 800bp including the ATG and signal sequence (Fig. 1Ai) and replace these with an attP recombination site. The resulting *Drosophila* stock facilitates simple insertion of modified *sog* sequences, such as point mutants (Fig. 2A), for expression under the endogenous *sog* promoter. In addition, regulatory elements located within *sog* introns, for instance the *sog* primary enhancer, remain intact (Markstein et al., 2002). The *white* gene was included in the HDR template and used as a marker to identify successful CRISPR events (Fig. 1Aii), before removal by Cre-Lox recombination (Fig. 1Aiii).

**Figure 1.**
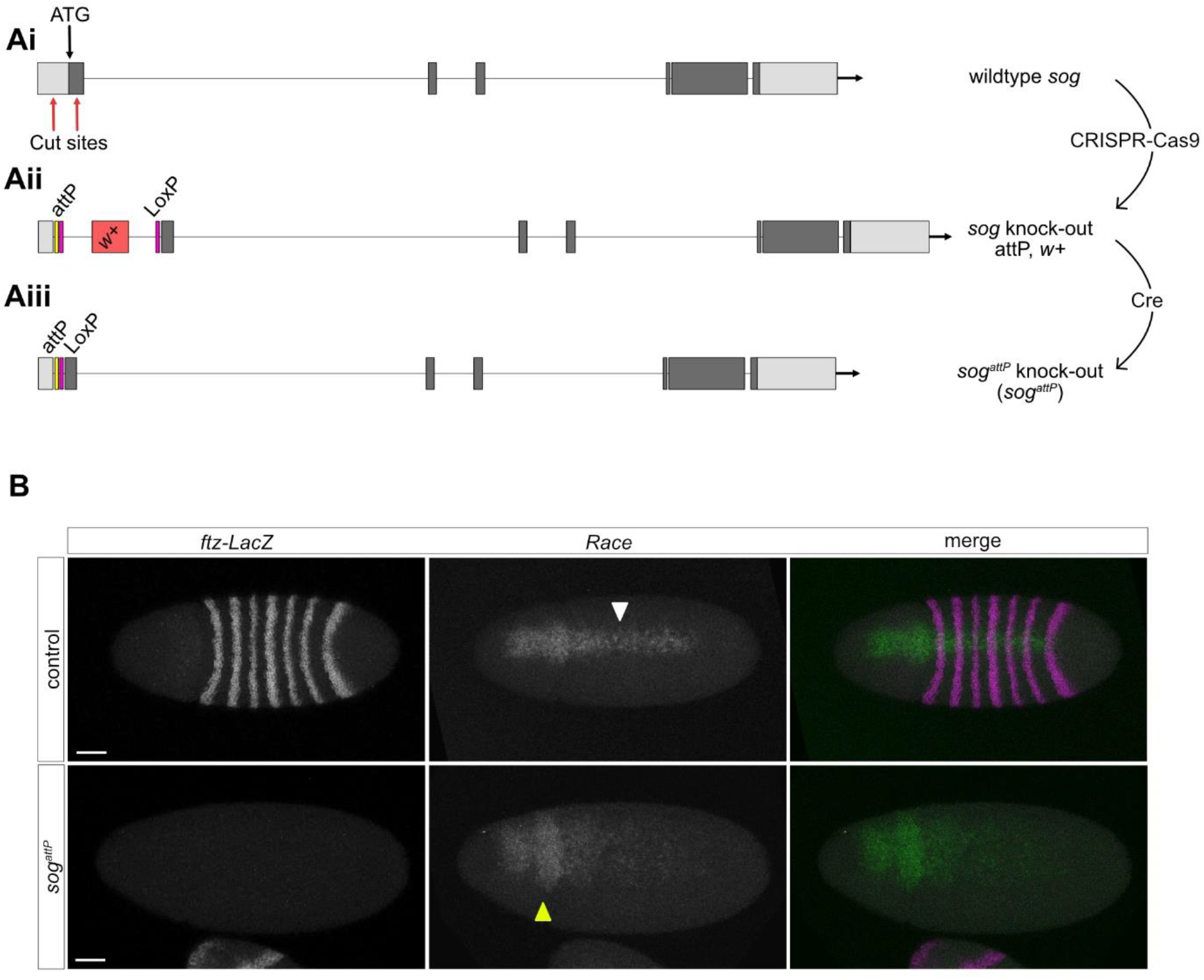
A CRISPR-Cas9 generated *sog* KO mutant. Ai) Cartoon shows the positions of the CRISPR-Cas9 cut sites (red arrows), which are located in the 5’UTR and downstream of the ATG and signal sequence in the first protein coding exon. Aii) CRISPR-Cas9 with HDR was used to insert an attP sequence, two LoxP sequences, and a *white+* marker gene. Aiii) Cre-Lox recombination removes the *white+* marker gene shown in Aii). The genome of the resulting fly line does not include the *sog* start codon or signal sequence, which are replaced with attP and LoxP sequences. (B) RNA *in situ* hybridisation for the BMP target gene *Race. Race* is expressed in heterozygous *sog*^*attP*^/FM7 *ftz-lacZ* embryos, the white arrowhead indicates *Race* expression in the presumptive amnioserosa (top panel). Heterozygotes are identified by *lacZ* expression from the FM7 *ftz-lacZ* balancer. *Race* expression is lost in the presumptive amnioserosa and the ‘head spots’ are broader (yellow arrowhead) in *sog*^*attP*^ embryos. Scale bars = 50 μm.

**Figure 2.**
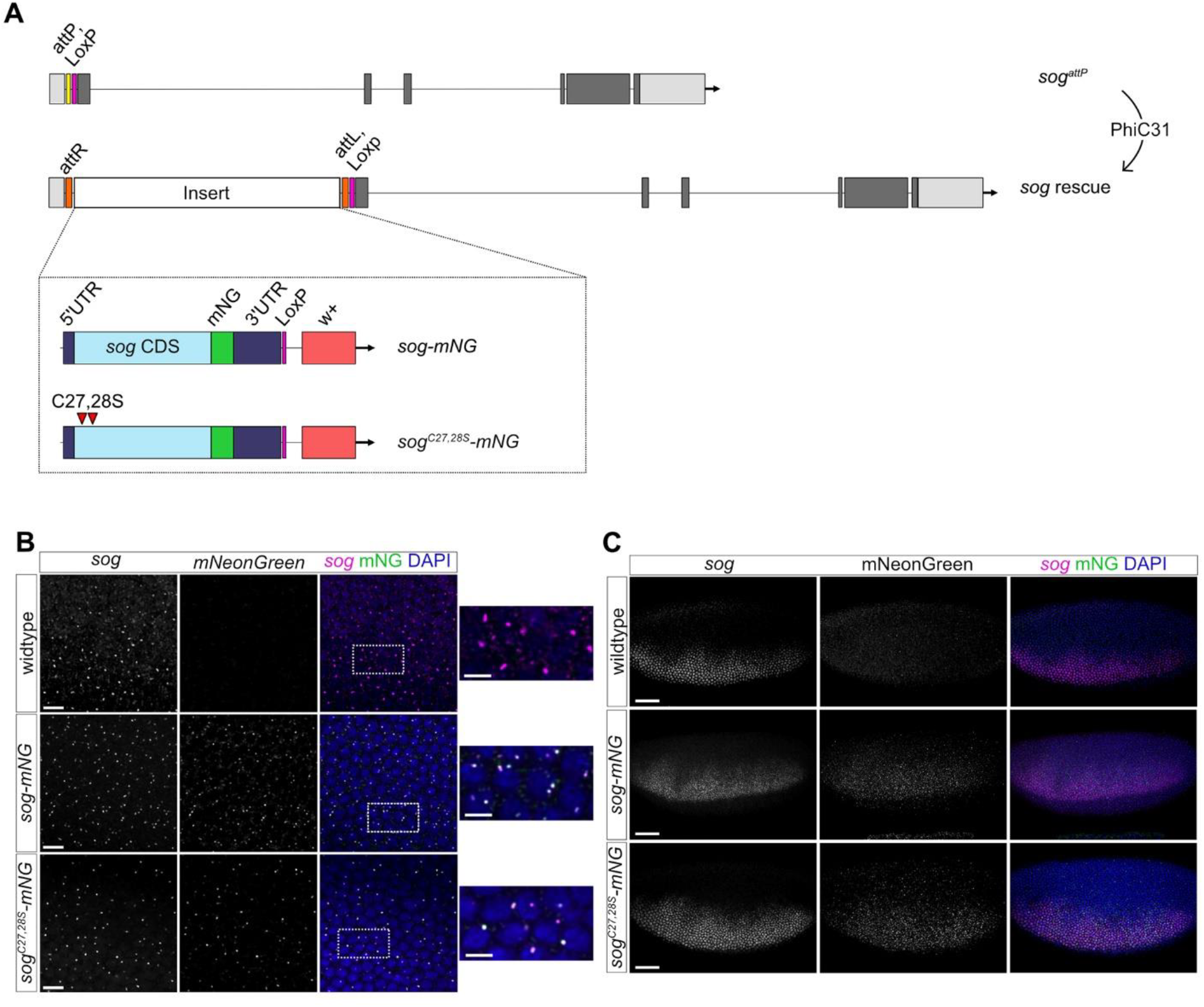
Insertion of specific *sog* coding sequences into the endogenous *sog* locus. (A) Specific *sog* coding sequences fused to a C-terminal mNG tag were inserted into the genome by phiC31 recombination at the inserted attP landing site of the *sog*^*attP*^ line. Wildtype Sog and a Cys27,28Ser mutant were reintegrated. (B) Embryos (nc14) of the indicated genotypes showing fluorescent RNA *in situ* hybridisation staining with *sog* and mNG probes (magenta and green, respectively). Scale bar = 10 µm. Expanded views of the areas outlined in the merged images are shown, scale bar = 5 μm. Colocalisation of *sog* (magenta) and mNG (green) transcription sites is indicated by white spots in the merged images. (C) *sog* smFISH (magenta) and mNG immunostaining (green) of the indicated nc14 embryos. A merged image with DAPI staining (blue) is shown, with single images of the *sog* and mNG channels for clarity. Scale bar = 50 μm.

The resulting *sog*^*attP*^ stock is maintained with an X-chromosomal balancer, and the absence of non-balancer males in the stock is consistent with a loss of Sog function. In addition, insertion of the attP recombination sequence at the *sog* locus was confirmed by sequencing. We used single molecule FISH (smFISH) to quantitate the amount of transcription from the *sog*^*attP*^ locus. The smFISH probes detect both *sog* transcription and mRNAs in early nuclear cycle (nc) 14 wildtype and *sog*^*attP*^ embryos (Fig. S1). To estimate *sog* mRNA number/cell, we assigned *sog* mRNAs to the closest nucleus as these embryos are only starting to cellularise. This analysis reveals that there is a ∼2.5-fold reduction in the peak numbers of *sog* mRNAs/cell in the *sog*^*attP*^ embryos (Fig. S1). The presence of *sog* mRNAs consistent with the deletion removing sequences downstream of the *sog* promoter, although the reduced mRNA number suggests that there is an effect on transcription and/or mRNA stability, potentially due to nonsense-mediated decay.

A scan for cryptic start codons at the modified *sog* locus identified one large and several smaller open reading frames (ORFs) (Fig. S2). Although the large ORF encodes Sog sequences starting within the first vWC domain, the signal sequence is absent. No cryptic signal sequence in this truncated Sog ORF was predicted using various software tools, e.g. Phobius webserver (Käll et al., 2004, 2007), that were able to predict endogenous Sog’s signal sequence/transmembrane domain (data not shown). As Sog is a secreted protein (François & Bier, 1995; Marqués et al., 1997), no Sog function is predicted following the deletion and attP insertion made in the *sog* locus.

To confirm loss of Sog function in *sog*^*attP*^ embryos, RNA fluorescence *in situ* hybridisation (FISH) was used to visualise expression of the peak BMP target gene *Race* (Fig. 1B). We also probed for *lacZ* mRNAs as *ftz-lacZ* is present on the balancer chromosome. The absence of *lacZ* expression indicates that the embryos are *sog*^*attP*^ males. In these *sog*^*attP*^ embryos *Race* expression is expanded in the anterior and lost in the presumptive amnioserosa (Fig. 1B), consistent with that described in *sog* null embryos (Ashe & Levine, 1999). These data, and the lethality of *sog*^*attP*^ males, support successful removal of the *sog* translation start site and its replacement with an attP recombination sequence to generate a *sog*^*attP*^ knockout (KO) allele.

### Reintegration of transgenes at the *sog* locus

The presence of an attP landing site at the *sog* locus facilitates targeted insertion of specific *sog* coding sequences into the genome (Fig. 2A). A previous study proposed that palmitoylation at two cysteines, at positions 27 and 28, is important for Sog secretion and stability of a membrane bound form of Sog (Kang & Bier, 2010). To test how the disruption of palmitoylation affects Sog function *in vivo* we used the *sog*^*attP*^ line we generated to integrate a *sog* cDNA in which Cys27 and 28 are mutated to Ser. Wildtype and palmitoylation mutant versions of the *sog* cDNA, to which a C-terminal mNeonGreen (mNG) tag was added (referred to as *sog-mNG* and *sog*^*C27,28S*^*-mNG*, respectively), were integrated into the endogenous locus (Fig. 2A). In total, ∼12.6 kb of DNA was inserted at the *sog* locus, including the *sog* CDS, *white+* marker, and LoxP sites (Fig. S3). Although endogenous *sog* sequences remain downstream of the integration site, cryptic initiation within the reintegration sequences and readthrough is not predicted to result in a Sog ORF longer than the truncated one described above. Therefore, if transcription of the remaining endogenous *sog* locus occurs due to cryptic initiation, the mRNA is only predicted to encode a truncated Sog ORF lacking a signal sequence (Fig. S2). This truncated Sog lacks activity based on the phenotype of the *sog*^*attP*^ embryos and lethality of the *sog*^*attP*^ males, as described above.

Both male and female flies carrying only the reintegrated *sog-mNG* and *sog*^*C27,28S*^*-mNG* sequences are viable (see later). Transcription of the integrated *sog* sequences in lateral stripes in the embryos was confirmed by FISH using *sog* and *mNG* probes (Fig. 2B). In this experiment, the control embryos carry an unmodified *sog* locus so a signal is only detected with the *sog* probe (Fig. 2B). However, both the *sog* and *mNG* probes detect co-localised signals in the *sog-mNG* and *sog*^*C27,28S*^*-mNG* embryos, as expected for transcription of the reintegrated sequences. We next used smFISH to test for any differences in *sog* expression between *sog-mNG* and *sog*^*C27,28S*^*-mNG* early stage 6 embryos compared to wildtype (Fig. S4A). We found no significant difference in the number of *sog* expressing cells between embryos of these genotypes (Fig. S4B). We were unable to quantitate absolute *sog* mRNA numbers at this stage due to their clustering. However, quantitation based on fluorescence intensity in equivalent areas of the expression domain suggests that there are no significant differences in *sog* expression levels in the reintegration embryos (Fig. S4C). The same result was obtained by analysing the fluorescence intensity along a line through the whole expression domain (data not shown). Finally, mNG immunostaining in combination with *sog* smFISH showed accumulation of Sog-mNG and Sog^C27,28S^-mNG protein (Fig. 2C).

### Cysteines 27 and 28, putative palmitoylation targets, are not essential for Sog function

As a fly stock homozygous for *sog*^*C27,28S*^*-mNG* was successfully established and maintained, cysteines 27 and 28 are not essential for Sog function. Due to difficulties associated with detecting palmitoylation *in vivo* we were unable to directly compare palmitoylation levels of wildtype and the mutant Sog. However, as the SogC27,28S mutant was less able to inhibit BMP activity in a tissue culture assay (Kang & Bier, 2010), we investigated whether these mutations reduce viability *in vivo*. To test this, the survival of embryos to pupal and adult life stages was quantified (Fig. S5A). *sog-mNG* or *sog*^*C27,28S*^*-mNG* embryos were raised at 25°C and the number of pupae and eclosed adults counted. The proportion of pupae and adults show some lethality at each of these stages for both the *sog-mNG* or *sog*^*C27,28S*^*-mNG* lines. This could be due to the presence of the mNG tag or differences in the reintegration locus compared to wildtype (see Discussion). Despite a trend towards lower survival rates for the *sog*^*C27,28S*^*-mNG* allele, there is no significant difference between the number of *sog*^*C27,28S*^*-mNG* and *sog-mNG* embryos that developed into pupae and successfully eclosed as adults.

To further test the functionality of the *sog-mNG* and *sog*^*C27,28S*^*-mNG* sequences, the extent to which these alleles can rescue the *sog*^*S6*^ loss-of-function allele or the *sog*^*attP*^ KO allele was assayed. *sog-mNG* and *sog*^*C27,28S*^*-mNG* males were crossed to *sog*^*S6*^/FM7 or *sog*^*attP*^/FM7 females (Fig. S5Bi), and the numbers of female offspring with either the *sog-mNG* or *sog*^*C27,28S*^*-mNG* allele versus the FM7 balancer were counted. No significant difference in the ability of the *sog-mNG* or *sog*^*C27,28S*^*-mNG* alleles to rescue either *sog* mutant allele relative to wildtype was observed (Fig. S5Bii). Although the different viability assays appear to have different sensitivities (Fig. S5A, B), together the data are consistent with the SogC27,28S mutations having only a very minor effect, if any, on Sog function.

### BMP signalling readouts in *sog-mNG* and *sogC27,28S-mNG* embryos

Although *sog-mNG* and *sogC27,28S-mNG* flies are viable, we investigated whether there are minor effects on Dpp gradient formation and interpretation. Sog functions in the early *Drosophila* embryo to concentrate BMP ligands dorsally, resulting in a stripe of the activated pMad transducer at the dorsal midline (Montanari et al., 2022). Therefore, pMad distribution was visualised in early stage 6 embryos by immunostaining and the width of the pMad stripe was measured at 50% embryo length (Fig. 3A & B). The pMad stripes in *sog-mNG* and *sog*^*C27*.*28S*^*-mNG* embryos are generally broader than those in wild-type control embryos, however these differences are not significant (Fig. 3B).

**Figure 3.**
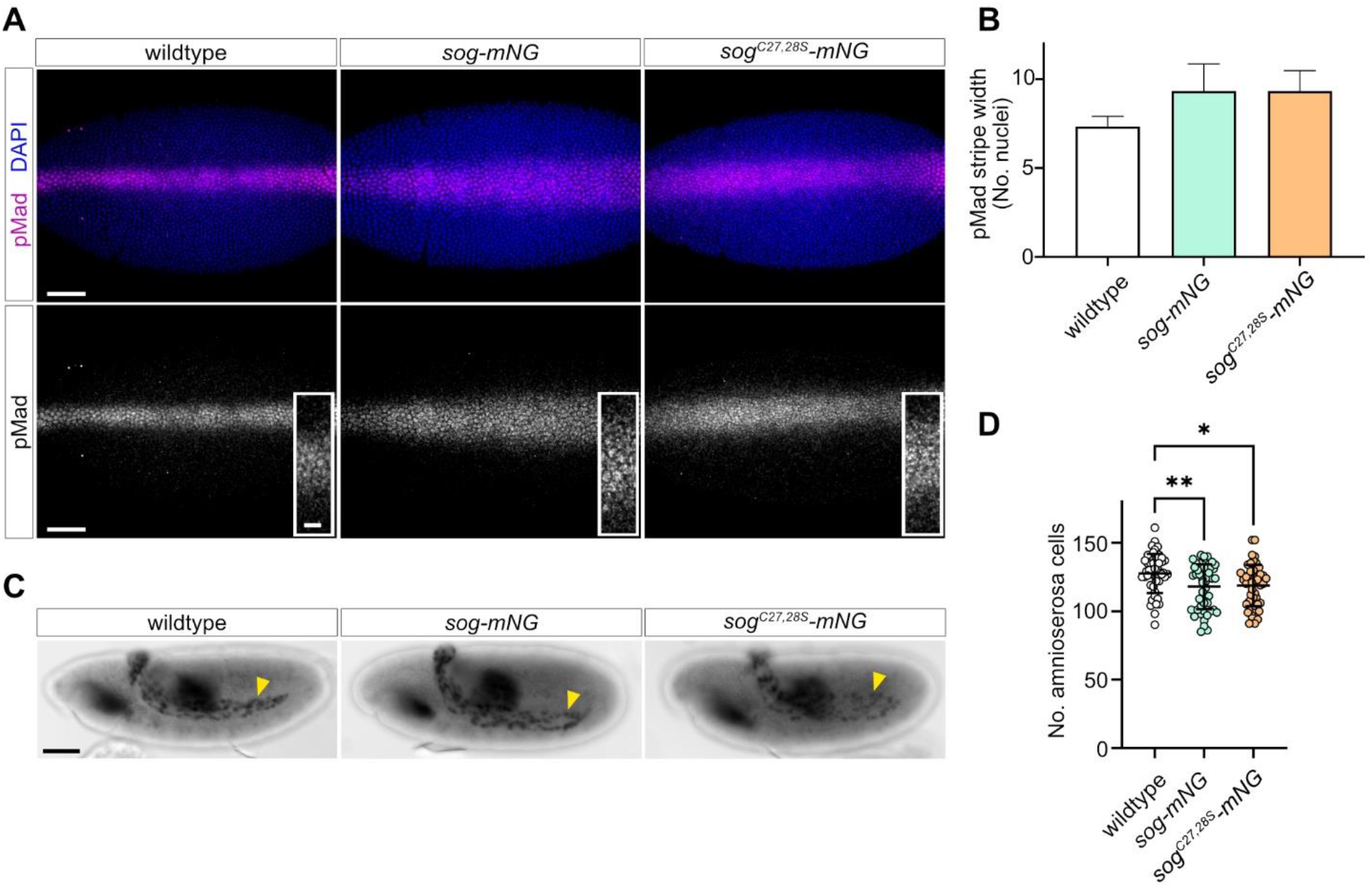
BMP signalling readouts in *sog-mNG* and *sogC27,28S-mNG* embryos. (A) pMad immunostaining in early stage 6 wildtype control, *sog-mNG*, and *sog*^*C27,28S*^*-mNG* embryos, scale bar = 50 μm. Insets show a higher magnification view of the central region of each pMad stripe, scale bar = 10 μm. (B) Mean pMad width (in number of nuclei) at ∼50% embryo length was measured for each embryo. No significant differences in mean pMad width were found by one-way ANOVA (with Tukey’s multiple comparisons test, P>0.05). For each genotype, n=3. Error bars represent mean + SD. C) Hnt immunostaining of stage 11 embryos shows amnioserosa cells (arrowheads) and midgut staining. Scale bar = 50 μm. D) Quantification of amnioserosa cell numbers in embryos of each genotype is shown. *sog-mNG* and *sog*^*C27,28S*^*-mNG* embryos have significantly fewer amnioserosa cells than controls (One-way ANOVA, Tukey’s multiple comparisons test, *sog-mNG* versus control P=0.006, *sog*^*C27,28S*^*-mNG* versus control P = 0.012), n = 50.

Peak BMP/pMad signalling specifies amnioserosa cell fate. Therefore, to test whether subtle differences in pMad stripe width affected amnioserosa specification, embryos were stained for the amnioserosa cell marker Hindsight (Hnt, Fig. 3C). Both *sog-mNG* and *sog*^*C27,28S*^*-mNG* embryos have a small but significant reduction in the number amnioserosa cells compared to wild-type embryos. However, the *sog-mNG* and *sog*^*C27,28S*^*-mNG* embryos have a similar reduction in the number of amnioserosa cells, suggesting that the C27,28S mutations do not affect embryonic BMP-dependent cell fate decisions. Together, these data suggest that BMP signal reception is marginally affected in both *sog-mNG* and *sog*^*C27,28S*^*-mNG* embryos, however this level of disruption is tolerated during development.

### Quantitative analysis of BMP target gene expression

As the number of amnioserosa cells was reduced in *sog-mNG* and *sog*^*C27,28S*^*-mNG* embryos, we used smFISH and quantitative analysis to assess effects on transcription of BMP target genes. smFISH was performed for the BMP target genes *Race* and *u-shaped* (*ush*), which respond to peak and intermediate levels of BMP signalling, respectively (Fig. 4A, B) (Ashe et al., 2000). Both *sog-mNG* and *sog*^*C27,28S*^*-mNG* embryos show similar *ush* expression patterns to wildtype (unedited) embryos, indicating that there is a BMP gradient, consistent with Sog function (Ashe et al., 2000). However, while the number of mature *ush* mRNAs in *sog-mNG* embryos is similar to that in wildtype embryos, *sog*^*C27,28S*^*-mNG* embryos have, on average, around half the number of *ush* mRNAs (Fig. 4C).

**Figure 4.**
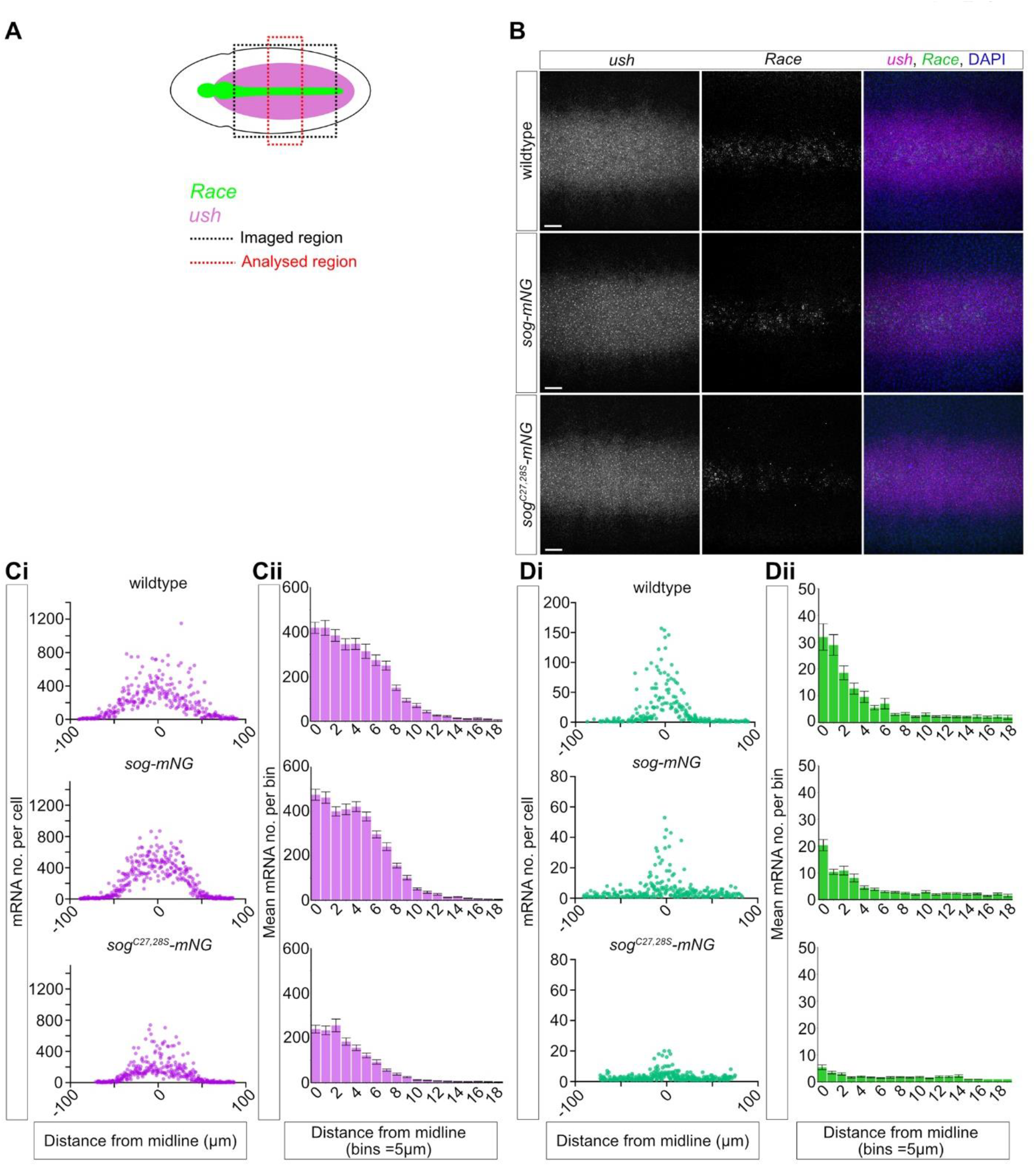
*sog*^*C27,28S*^*-mNG* embryos have disrupted BMP target gene expression. (A) Cartoon of a stage 6 *Drosophila* embryo showing the *Race* and *ush* expression patterns in green and magenta, respectively. The black box indicates the imaged region of the embryo, the red box represents the portion of the images used for analysis. (B) smFISH detection of *ush* (magenta) and *Race* (green) mRNAs in wildtype, *sog-mNG*, and *sog*^*C27,28S*^*-mNG* stage 6 embryos. Scale bars = 20 μm. (Ci) The numbers of *ush* transcripts per cell were quantified and plotted against the distance from the dorsal midline (μm, midline = 0). Analysis was performed for three embryos of each genotype, the results for one representative embryo of each genotype are shown here (see Fig. S6 for the other embryos). (Cii) Counts of *ush* mRNAs per cell for the three replicates of each genotype were combined and binned according to distance from the midline (bins = 5 μm, each bin approximately corresponds to a nuclear width). Error bars represent mean ± SEM. (Di, ii) Data shown are as for (Ci,ii), but for *Race*. Note the change in the axis scale for the control embryo in (Di). Error bars represent mean ± SEM.

*Race* expression levels in *sog-mNG* embryos are, in general, lower than in controls, but *Race* is restricted to the dorsal midline as in control embryos (Fig. 4D, Fig. S6). The mean number of *Race* mRNAs across the 3 biological repeat embryos is slightly lower in *sog-mNG* relative to control embryos, whereas there is an even greater reduction in *Race* expression in *sog*^*C27,28S*^*-mNG* embryos (Fig. 4D). In addition, the levels of *Race* expression observed in *sog*^*C27,28S*^*-mNG* embryos show more variation: while one *sog*^*C27,28S*^*-mNG* embryo has a weak stripe of *Race* expression along the dorsal midline (Fig. 4Di), it is almost absent in the others (Fig.S6). As *Race* expression in *sog*^*C27,28S*^*-mNG* embryos is weaker than in *sog-mNG* embryos, this suggests that Sog^C27,28S^-mNG may be less able to promote peak BMP signalling. Overall, this highly sensitive assay of BMP target gene transcription identifies subtle deficiencies in the responses, particularly with Sog^C27,28S^-mNG, even though these do not have major effects on viability.

### The Sog C27,28S mutant shows weakly penetrant PCV patterning defects

Sog also regulates BMP signalling during pupal wing vein patterning, including PCV patterning. Therefore, we used this as an alternative developmental context to test whether the requirement for palmitoylation of Sog may be context dependent. The wings of adult female flies, raised at either 18°C or 25°C, were examined for defects in PCV specification and patterning (Fig. 5). A low proportion of *sog*^*C27,28S*^*-mNG* wings displayed a mutant PCV phenotype: a small extension to the distal side of the PCV (Fig. 5A). Ectopic PCV development was observed in a slightly higher proportion of flies that developed at 18°C compared to 25°C, suggesting that the phenotype is exacerbated by mild cold temperature stress (Fig. 5B, C). Disruption to the PCV only in *sog*^*C27,28S*^*-mNG* wings at 18°C suggests that mutation of cysteines 27 and 28 has mildly impacted Sog function or levels, resulting in reduced BMP signal refinement during PCV patterning (Antson et al., 2022) (see Discussion).

**Figure 5.**
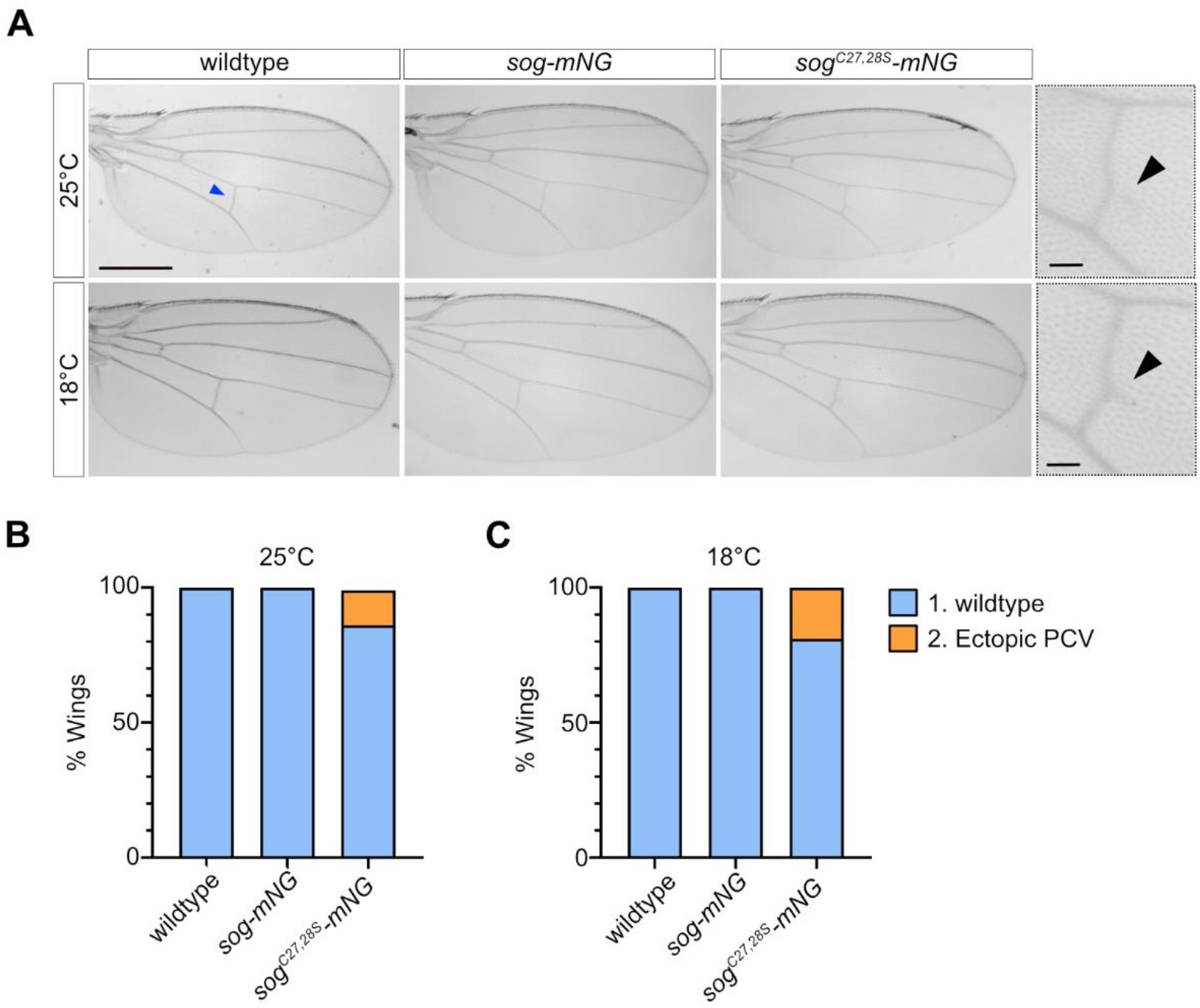
The Sog C27,28S mutant shows weakly penetrant PCV patterning defects. (A) Wings from female control, *sog-mNG*, and *sog*^*C27,28S*^*-mNG* adults raised at 18°C and 25°C (scale bar = 500μm). The wing PCV (blue arrowhead) was classed as wildtype or ectopic PCV. Higher magnification views of *sog*^*C27,28S*^*-mNG* wings with a mild ectopic PCV phenotype are shown adjacent to images of the whole wing (scale bar = 50μm). The black arrowhead indicates ectopic extension to the distal side of the PCV. (B-C) Percentage of wings in each phenotypic class at 25°C (B) and 18°C (C). Samples sizes for flies raised at 25°C are as follows: control n=56, *sog-mNG* n=66, *sog*^*C27,28S*^*-mNG* n=67. Sample sizes for flies raised at 18°C: control n=64, *sog-mNG* n=65, *sog*^*C27*.*28S*^*-mNG* n=67. Fisher’s exact test (two-sided) finds a significant association between genotype and PCV phenotype in flies raised at 25°C (p<0.01) and ta 18°C (p<0.01). Statistical tests were performed on raw data.

### Sog has a curved shape

The data presented above demonstrate the utility of our *sog*^*attP*^ line for testing and elucidating the effect of specific *sog* mutations. One limitation for targeted mutagenesis of *sog*, however, is the absence of structural information. Therefore, to investigate Sog structure, we purified Sog (with C-terminal His and V5 tags) from the conditioned media of a stable, Sog-expressing, HEK293 EBNA cell line by affinity purification followed by two rounds of size exclusion chromatography (SEC) (Fig.6A, Fig. S7). Sog purity was assessed by SDS-PAGE and western blot analysis where, after an initial round of SEC, a prominent doublet band was typically observed (Fig. S7C). This doublet band is likely to represent full-length Sog and Sog lacking the N terminal vWC1 domain due to Tld cleavage, as the cleavage product is detected by a His antibody (Fig. S7C) and expression of Chordin in the same cell line results in co-purification of a Tolloid cleavage product lacking the vWC1 domain (Troilo et al., 2014). A second round of SEC was included but could not completely separate the lower molecular weight species (Fig 6A and B). Negative stain transmission electron microscopy (TEM) was used to investigate the 3-dimensional structure of the purified Sog protein. During single particle analysis, 2D classification aligned Sog particle images and produced 2D class averages that were used to generate and refine a Sog 3D model (Fig. 6C, D). The final 3D reconstruction, with an estimated resolution of 22.8 Å (Fig. 6D, E), reveals that Sog has an asymmetric, curved shape and dimensions of 13.6 nm X 9.9 nm X 9.2 nm (Fig. 6D). Due to the similarity in size, the Sog cleavage product cannot be separated from the full-length protein during image analysis, therefore the cleavage product will also have a contribution to the 3D model.

**Figure 6.**
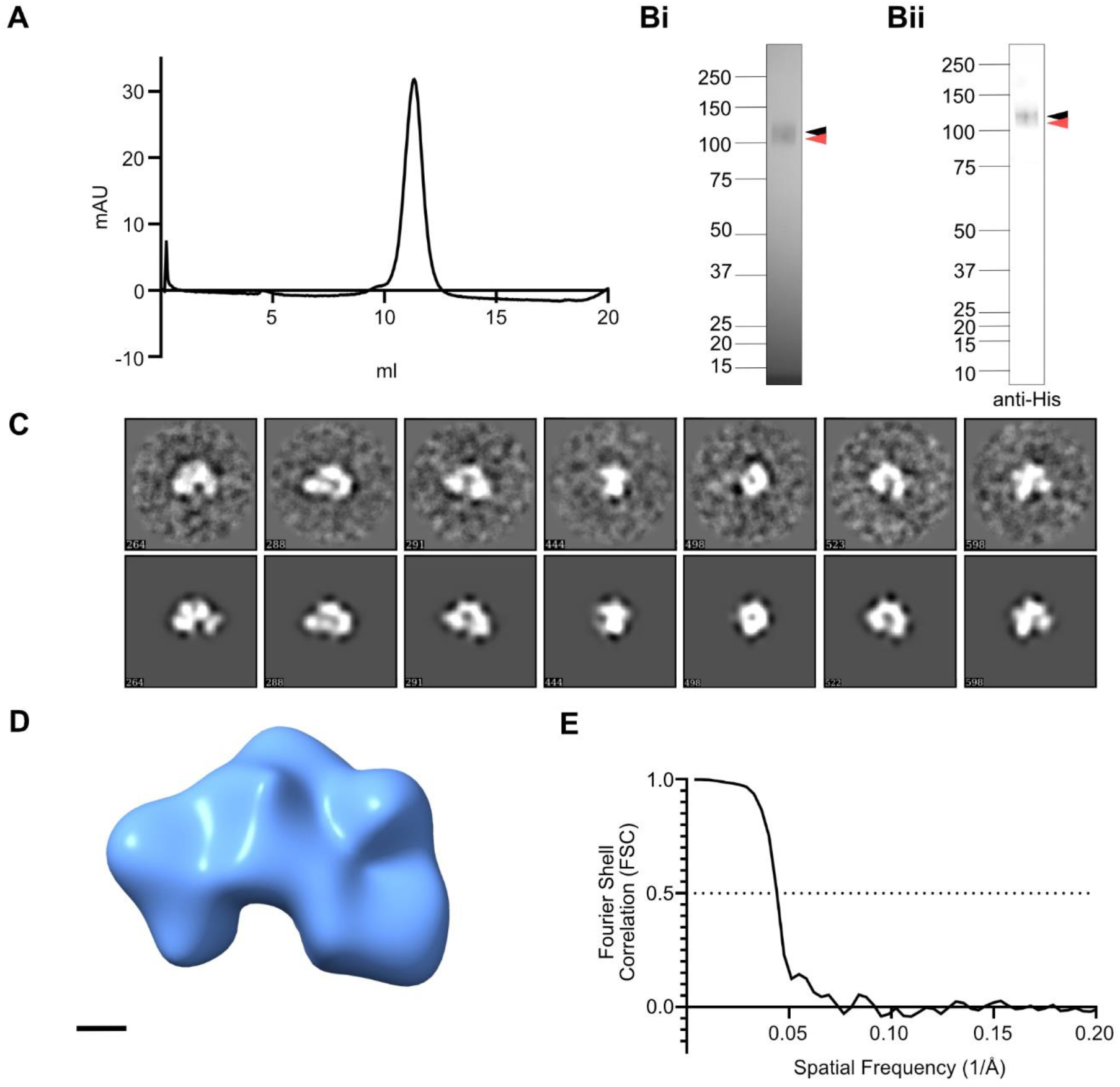
Sog has a curved shape. (A) SEC trace for Sog purification. The Sog sample was subject to 2 rounds of SEC to minimise the contribution of a Tld cleavage product. The SEC trace for the second SEC round is presented here, see also Fig. S7 for the first SEC purification. (B) Reduced SDS-PAGE gel (i), and anti-His western blot (ii), corresponding to the SEC trace in (A), showing purified Sog. Full length Sog is indicated by a black arrowhead. Full length Sog co-purifies with an Sog N-terminal cleavage product (red arrowhead). (C) A selection of 2D class averages (top panel) generated during the final refinement and corresponding re-projections (bottom panel) of the Sog model shown in (D). Numbers in the bottom left of the boxes are reference numbers arbitrarily assigned to classes and projections by the analysis software. Box size = 36 nm. (D) Final 3D model of Sog. Scale bar = 2 nm. (E) Fourier Shell Correlation (FSC) calculations were used in Eman2.2 (Tang et al., 2007) to estimate the model resolution using a cut off of 0.5.

### Comparing AlphaFold predictions to experimental data

Next we probed the AlphaFold protein structure database, a recently developed resource based on the machine learning prediction of protein structures to atomic resolution (Jumper et al., 2021; Varadi et al., 2022), to analyse the predicted Sog structure and investigate how it fits within the Sog 3D reconstruction described above. For reference, the domain organisation of Sog is shown in Fig. 7A. Given the presence of the hydrophobic N-terminally located Sog transmembrane domain/signal peptide, it is likely that much of the Sog N-terminus is cleaved prior to secretion into the perivitelline space. The Sog N terminus, up to H80, has therefore been removed from the AlphaFold model, at a site between R79 and H80 previously identified as a putative cleavage target for separation of a hydrophobic N-terminal signal sequence/transmembrane domain from mature extracellular Sog (Shimmi & O’Connor, 2003). AlphaFold generates a pLDDT score, which is a useful metric from which to infer confidence in the local predicted protein structure. AlphaFold predicts the folds of each Sog vWC domain, with the majority of residues predicted with a high pLDDT score of 70-90 (Fig. 7B). Furthermore, AlphaFold predicts a novel fold for each of the four CHRD specific domains (Fig.7A) with high confidence (pLDDT >90 & 70-90) (Fig. 7B). Residues linking vWC domains to other vWC or CHRD domains, however, are predicted with low pLDDT scores (50-70 & <50), indicating potentially disordered and/or flexible regions.

**Figure 7.**
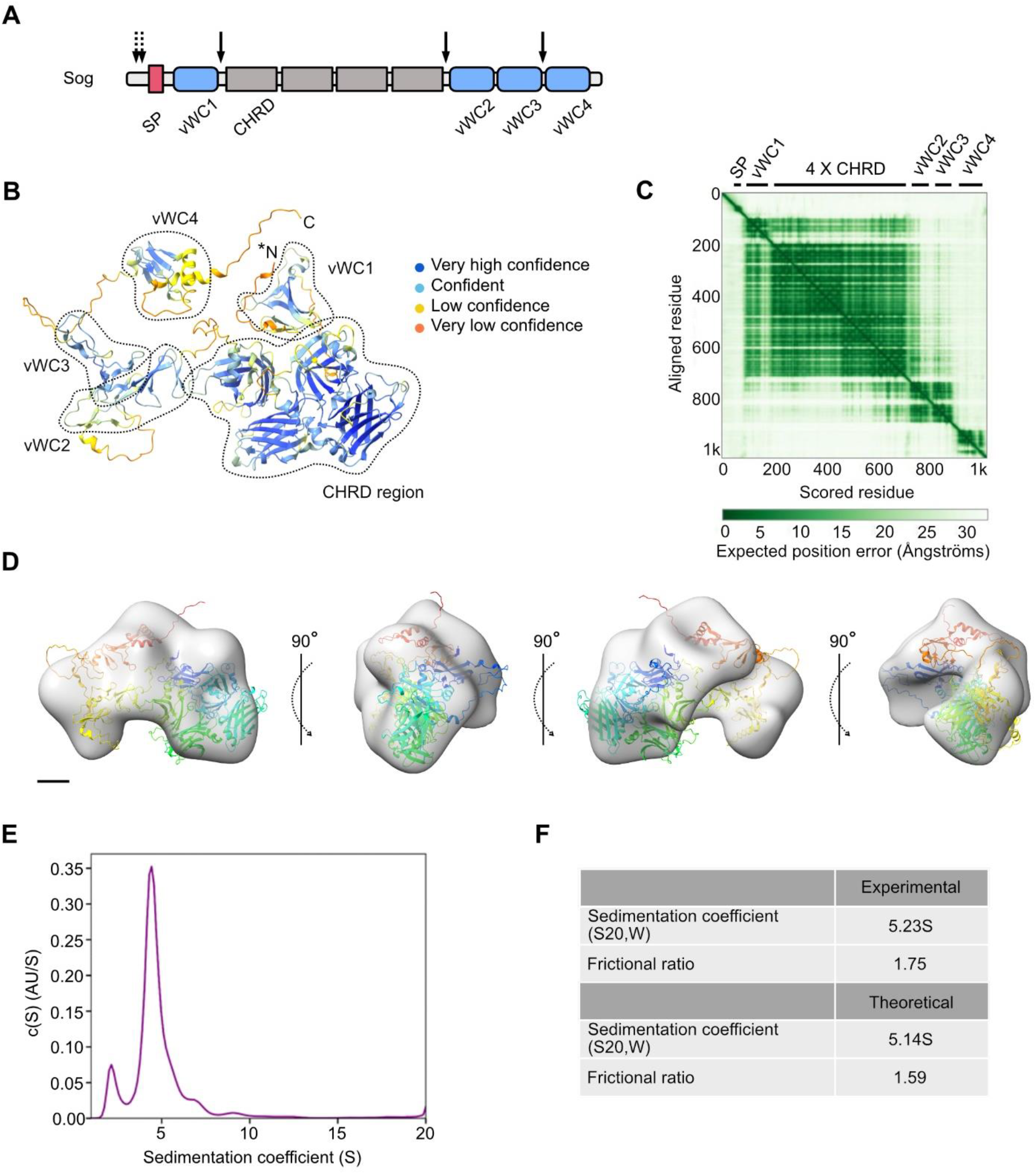
Sog negative stain EM model is consistent with the AlphaFold prediction of Sog structure. (A) Sog comprises four vWC domains (blue) and four CHRD domains (grey). The N-terminal Sog hydrophobic domain is predicted to be a transmembrane domain (TMD) or signal peptide (red, SP). Solid arrows indicate Tld cleavage sites, dashed arrows indicate the location of the putative palmitoylation sites cysteines 27 and 28. (B) The prediction of the Sog structure by AlphaFold (Jumper et al., 2021; Varadi et al., 2022). N- and C-termini are indicated, the model shown here has been N-terminally truncated between R79 and H80 (N*). Sog vWC and CHRD domains are circled (figure adapted from AlphaFold entry Q24025). The predicted model is coloured according to per residue confidence (pLDDT) score. A score of 100 indicates maximum confidence. pLDDT scores are as follows: dark blue, > 90; blue, 70-90; yellow, 50-70; orange, <50. (C) Predicted aligned error (PAE) (Figure from AlphaFold entry Q24025). Domain boundaries are indicated by black bars along the ‘scored residue’ (X-) axis. (D) AlphaFold Sog model, N-terminally truncated after the hydrophobic domain at the predicted cleavage site, fits within the Sog EM model volume. Scale bar = 2 nm. (E) Sedimentation coefficient distribution produced by c(S) analysis of AUC data. After accounting for the sample buffer, the sedimentation coefficient is calculated to be 5.23 S (S20,W). (F) A comparison of experimental and theoretical sedimentation coefficients and frictional ratios. Theoretical values were calculated with US-SOMO software (Brookes et al., 2010; Spotorno et al., 1997) for the predicted Sog structure by AlphaFold.

A Predicted Aligned Error (PAE) score is also calculated for each residue pair by AlphaFold (Fig.7C) (Jumper et al., 2021; Varadi et al., 2022). This score is a measure of the confidence with which the positions of amino acid pairs are predicted, thereby indicating the confidence of relative domain positions. The relative positions of each CHRD domain, vWC1 and the 4x CHRD region, and vWC2 and vWC3 are predicted with high confidence. Other inter-domain distances, for instance between vWC3 and vWC4, are predicted with lower confidence, suggesting some flexibility in the full-length protein. The Tld cleavage sites in both Sog and its ortholog Chordin are located within these flexible interdomain regions (Fig.7A & Fig. S8) (see Discussion).

The asymmetric nature of the AlphaFold model, with the bulky 4x CHRD region on one side (Fig. 7B), resembles the Sog 3D negative stain EM model (Fig. 6D). Indeed, overlay of the AlphaFold model with the Sog 3D EM volume, using ChimeraX (Pettersen et al., 2021), demonstrates how the 4x CHRD region might sit within the larger arm of the Sog EM model, as well as the arrangement of the other domains within the density (Fig. 7D). For further insight into the hydrodynamic properties of Sog, purified Sog protein was subject to sedimentation velocity analytical ultracentrifugation (AUC) (Fig. 7E). c(S) analysis calculated a sedimentation coefficient of 5.23 (S20,W), and a frictional ratio of 1.75, reflecting the relatively large size of Sog and indicating an elongated shape (Fig. 7F). To further test the level of agreement between the Sog AlphaFold structure and purified Sog, the sedimentation coefficient and frictional ratio of the N-terminally cleaved Sog AlphaFold model was predicted with US-SOMO (Spotorno *et al*., 1997; Rai *et al*., 2005; Brookes, *et al*., 2010). A sedimentation coefficient of 5.14 (S20,W) and frictional ratio of 1.59 were predicted for the Sog AlphaFold model (Fig. 7F), lending further confidence to the accuracy of this predicted atomic structure. Together, similarities in shape between the AlphaFold Sog prediction and the Sog EM model, and between experimentally and theoretically derived hydrodynamic parameter values, suggest an overall domain organisation that provides a framework for future studies.

## Discussion

In this study, we have generated a *sog* knockout line that allows simple reintegration of altered *sog* cDNAs. While the Sog 3D model that we have described will allow targeted mutagenesis in future studies, here we showed the utility of our *sog*^*attP*^ KO stock by using it to investigate the effect of mutating two residues implicated in Sog palmitoylation. Palmitoylation at cysteines 27 and 28 has previously been suggested to play an important role in membrane targeting and Sog secretion (Kang & Bier, 2010). However, our data show that mutation of these two residues to serine *in vivo* resulted in viable flies, consistent with these residues not being essential for Sog function.

Our data reveal that the wildtype *sog-mNG* reintegration embryos accumulate less *Race* mRNAs and have a minor reduction in the number of amnioserosa cells compared to wildtype embryos. As *Race* expression and amnioserosa fate are dependent on peak signalling, we speculate that there is a subtle defect in BMP gradient formation arising from slightly reduced shuttling of BMP heterodimers to the dorsal midline in the *sog-mNG* embryos. Although no significant difference in the width of the pMad stripe in the reintegration embryos was observed, the reduced *Race* mRNA numbers suggest that there is a minor defect in pMad levels. Co-staining of wildtype and *sog-mNG* or *sog*^*C27,28S*^*-mNG* embryos with the pMad antibody, along with sophisticated quantitation of the staining intensities (Gavin-Smyth et al., 2013; Umulis et al., 2010) will allow this to be addressed in future studies.

There are various potential explanations for the mildly reduced peak outputs in the *sog-mNG* reintegration embryos. Fusion of the C-terminal mNeonGreen tag to Sog may have slightly reduced its function. It is also possible that we have perturbed the timing of expression as we reintegrated the *sog* cDNA. As the cDNA is much shorter than the sequences present in the endogenous locus that contains long introns, there is potentially more rapid accumulation of *sog* mRNAs, which may be particularly important in the short nc13 (Sandler et al., 2018). However, we did not detect any global effects on *sog* expression levels relative to wildtype embryos at nc14, which may be expected if there is earlier accumulation of full length *sog* mRNAs. Introduction of the *sog* cDNA in the reintegration embryos also prevents expression of a truncated *sog* transcript that includes intron-derived sequence. The encoded short Sog protein has been shown to suppress early BMP signalling and prevent ectopic target gene expression during nc13 (Sandler et al., 2018). However, as we observe a reduction in *Race* mRNAs at nc14, this phenotype does not seem compatible with a loss of Short Sog expression. Reintegration also introduces additional sequences including an attR scar in the 5’UTR, which could affect translational regulation of the *sog* mRNA. In addition, the endogenous *sog* locus sequences (starting at the end of the CRISPR deletion) are present downstream of the reintegration sequences. While we find no evidence that the remnants of the endogenous locus can encode a secreted functional Sog protein, these sequences can be removed in future studies by Cre recombination.

Using the *sog-mNG* flies as controls, we identified two minor phenotypes associated with *sog*^*C27,28S*^*-mNG* embryos and wings. There is a reduction in both *Race* and *ush* expression in the *sog*^*C27,28S*^*-mNG* embryos, suggesting that there are slightly lower levels of the pMad activator. We speculate that this is due to a further minor reduction in BMP shuttling by the mutant Sog, due to a slight decrease in either its levels or activity, resulting in a lower concentration of pMad at the dorsal midline. Although *sog*^*C27,28S*^*-mNG* embryos have ∼4-fold reduction in *Race* mRNA numbers relative to the *sog-mNG* embryos, there is no difference in the number of amnioserosa cells specified. As *Race* expression is exquisitely sensitive to reductions in peak Dpp/Scw signalling (Ashe & Levine, 1999; Rusch & Levine, 1997), it is possible that other peak BMP targets show a more modest reduction in mRNA numbers. Consistent with this, the intermediate target gene *ush* is only reduced ∼2-fold in the *sog*^*C27,28S*^*-mNG* embryos. We also observed a weakly penetrant ectopic PCV phenotype in *sog*^*C27,28S*^*-mNG* wings, as a minor proportion have a small extension to the distal side of the PCV. This phenotype is consistent with reduced pMad, potentially due to lower activity of the SogC27,28S mutant in promoting BMP transport to the PCV (Antson et al., 2022). While both the embryonic target gene and wing PCV phenotypes suggest slightly reduced pMad levels, overall these are very mild phenotypes and the flies with SogC27,28S are viable.

While our data suggest that C27 and C28 in Sog are not critical, previously it was shown that overexpression of the dHIP14 palmitoyl transferase in the anterior of the early embryo or wing reduced pMad and disrupted wing vein patterning, respectively, similar to the phenotypes associated with *sog* overexpression (Kang & Bier, 2010). It is possible that dHIP14 overexpression has pleiotropic effects. Recent evidence suggests that the inhibitory Smad Dad is palmitoylated, which is important for its function (Li et al., 2017). While *dad* is not expressed in the early embryo, perhaps effects on Dad palmitoylation contribute to the wing phenotypes observed on dHIP14 overexpression.

As the low levels of Sog *in vivo* prevented us from directly measuring its level of palmitoylation for the wildtype and C27,28S mutant, we cannot rule out another palmitoylation target site in Sog. A cys residue within the predicted TMD/SP is another putative target (Kang & Bier, 2010). However, conditioned media collected from cells expressing the Sog C27,28S mutant was less able to inhibit Dpp signalling compared to wild-type Sog in a tissue culture assay, consistent with reduced Sog secretion (Kang & Bier, 2010). This result suggests that the C27,28S mutation would be sufficient to reveal some defect in BMP signalling regulation *in vivo*, if Sog palmitoylation at these residues is important. Together, our data suggest that Sog secretion is much less dependent on C27, 28 and palmitoylation *in vivo* than in tissue culture cells.

Our 3D model of Sog constructed from negative stain EM data reveals a curved shape similar to the ‘horseshoe’ shape of its vertebrate ortholog Chordin. Due to the known BMP-Chordin vWC domain binding affinities, BMP dimers have been predicted to cooperatively bind Sog vWC1 and vWC4, and Chordin vWC1 and vWC3 domains (Larraín et al., 2000; Troilo et al., 2014; Zhang et al., 2007). In addition, Sog vWC1 and vWC4 have also been shown to interact with Col IV (Sawala et al., 2012). Sog and Chordin have therefore been predicted to adopt a curved conformation that would position the N- and C-termini in close proximity (Larraín et al., 2000; Sawala et al., 2012; Troilo et al., 2014). For Chordin, this ‘horseshoe-like’ shape is supported by a 3D reconstruction generated by single particle analysis of negative stain EM data (Troilo et al., 2014). The curved shape of the Sog EM and AlphaFold structures is therefore consistent with models of cooperative BMP binding, and simultaneous vWC1- and vWC4-Col IV interactions.

In comparison to Chordin, the model of Sog presented here shows Sog to be slightly more compact, with dimensions of 13.6 × 9.9 × 9.2nm versus 15 × 13 × 8nm of Chordin (Troilo et al., 2014). While Tld can only cleave Sog when it is bound to BMP, Chordin alone is processed by Tld (Marqués et al., 1997; Peluso et al., 2011). It has been suggested that a BMP induced conformational change in Sog is required for Sog cleavage by Tld (Marqués et al., 1997; Peluso et al., 2011; Serpe et al., 2005). The PAE scores for AlphaFold Sog suggest that it is less flexible than Chordin, which together with the more compact shape of Sog could contribute to the requirement for this conformational change. The lower confidence in the relative positions of C-terminal vWC domains of the AlphaFold model is consistent with a level of Sog flexibility, potentially facilitating a ligand dependent conformational change, and permitting Tld access to target residues. The predicted greater flexibility of Chordin interdomain regions, where Tld cleavage sites are located, could therefore reflect the absence of required co-factors for Tld-mediated Chordin processing. The dependency of Sog on BMP ligand binding for Tld cleavage appears to underpin the ‘shuttling’ function of Sog during *Drosophila* embryogenesis (Peluso et al., 2011). In contrast, a ‘source-sink’ model of BMP gradient formation is most likely to operate during vertebrate embryo dorsal-ventral patterning (Pomreinke et al., 2017; Tuazon et al., 2020; Zinski et al., 2017). Future studies probing how the structures of Sog and Chordin differ will illuminate how these proteins use the same interacting proteins but different mechanisms to generate a BMP gradient.

## Methods

### Sog purification

Lipofectamine 3000 reagent (Thermo Fisher Scientific) and Xfect (Takara Bio) transfection reagents were used to transfect HEK293 EBNA cells (Baldock lab stock, not recently authenticated) with pCep-Pu-Sog-V5-His to establish a stable cell line. pCep-Pu-Sog-V5-His cells were maintained in growth media (10% FBS, DMEM:F-12 Hams, 1% Penicillin-Streptomycin (P/S), 1% Glu) containing 1 μg/ml puromycin. For protein expression, cells were cultured in HYPERflasks (Corning) with expression media (DMEM:F-12 Hams, 1% P/S, 1% Glu, 50 mM L-Arginine). Collected conditioned media was stored at -20°C.

Recombinant Sog protein was isolated from collected conditioned media using affinity chromatography. 1 ml HisTrap Excel columns (Cytiva) were used to pull down Sog via the C-terminal His-tag. For this, 2 mM imidazole was added to filtered conditioned media, and a loading buffer was used (10 mM Tris, 800 mM NaCl, pH7.4) to equilibrate the His-trap column. Filtered conditioned media was pumped over the column at 4°C. The column was then washed with 20 column volumes (CV) loading buffer with 10 mM imidazole. Protein was eluted from the column with elution buffer (10 mM Tris, 800 mM NaCl, 500 mM imidazole, pH7.4).

Affinity chromatography fractions were subject to Size exclusion chromatography (SEC) for further purification with a Superdex 200 Increase 10/30 gel filtration column (Cytiva). The column was washed with 1.5 CV filtered and degassed Milli-Q water (Millipore), followed by 1.5 CVs SEC buffer (800 mM NaCl, 2.7 mM KCl, 10 mM Na2HPO4, 1.8 mM KH2PO4, pH 7.4) to equilibrate. Affinity chromatography fractions were passed across the column at 0.5 ml/min. Protein elution was monitored using UV absorbance (280 nm). Eluted 0.5 ml fractions were collected and screened by SDS-PAGE under reducing conditions with Instant Bluestain (Abcam), and by western blot with anti-His [clone AD1.1.10] (1:1000, R & D Systems Cat# MAB050, RRID: AB_357353) primary antibody and IRDye 800CW Donkey anti-Mouse (1:10,000, LI-COR Biosciences Cat# 926-32212, RRID: AB_621847Li-COR) secondary antibody. Purified Sog was stored at -80°C, before undergoing a second round of SEC (as above) to improve sample purity for production of negative stain EM grids. Collected fractions were screened by SDS-PAGE under reducing conditions with Colloidal Coomassie stain, and by western blot (as above).

### Negative stain electron microscopy

A 5 nm (approximately) layer of carbon was deposited onto a mica sheet using a Cressington coating system 308R. The carbon layer was floated onto the carbon side of the TEM grids which were left to dry overnight at room temperature. Carbon coated grids were glow discharged at 25 mA for 1 minute. After SEC, Sog (∼13 μg/ml) was adhered to carbon coated EM grids for 1 minute and stained with 2% uranyl acetate. The stain was wicked away with filter paper before drying. Negatively stained protein molecules were imaged in low dose mode on an FEI Tecnai 12 Biotwin Transmission Electron Microscope at 100 kV. Images were captured with a Gatan Orius SC1000 camera at 30,000 X magnification. Data collection parameters were as follows: Defocus = 0.5-1 μm; Exposure length = 0.6 s, pixel size = 2.8 Å.

Data collected from negative-stain EM grids were processed using Eman2.2 software (Tang et al., 2007). Particles were manually picked and processed with a box size of 128 pixels. Contrast Transfer Function (CTF) parameters were estimated in Eman2.2 and particle images underwent phase flipping to correct the CTF. A soft Gaussian mask, with an outer radius of 52 nm was applied to selected particles. 2D class averaging was performed to iteratively align and average the particles into 100 reference free class averages. Class averages were discarded based on particle contrast against background noise. Of the remaining classes, 17 were selected that represented different views of Sog to generate an initial model. The initial model underwent two iterative refinements, using a ‘gold standard’ refinement procedure to produce a final model. The Fourier shell correlation (FSC) of two separately refined half-maps was used to estimate the resolution of the final model, with the 0.5 threshold.

### AUC

A purified Sog sample (0.1 mg/ml) in the same buffer as used in SEC was characterized by sedimentation velocity AUC using a Beckman XL-A analytical ultracentrifuge with an An60Ti 4-hole rotor running at 45,000 rpm at 20°C. The sedimenting boundary was monitored at 230 nm for 300 scans. Data were analysed by continuous model-based distribution C(s) of Lamm equation solutions method using SEDFIT software (Schuck, 2000), and the resulting sedimentation coefficients were corrected to standard conditions using SEDNTERP software (Philo, 1997). For theoretical predictions of hydrodynamic properties for the predicted Sog structure by AlphaFold, US-SOMO software (Spotorno *et al*., 1997; Rai *et al*., 2005; Brookes, 2010; Brookes, *et al*., 2010; Brookes and Rocco, 2018) was used.

### Fly stocks

All stocks were grown and maintained on standard fly food media (yeast 50g/L, glucose 78g/L, maize 72g/L, agar 8g/L, 10% nipagen in EtOH 27mL/L and proprionic acid 3mL/L). The following fly lines were used in this study: *y*^*1*^ *w*^*67c23*^ (BDSC Stock 6599), *w*^*1118*^; *PBac{y[+mDint2] GFP[E*.*3xP3]=vas-Cas9}VK00027* (BDSC Stock 51324); *y*^*1*^ *w*^*67c23*^; *MKRS, P{ry+t7*.*2=hsFLP}86E/TM6B, P{w+mC=Crew}DH2, Tb1* (BDSC Stock 1501); *y*^*1*^ *sog*^*S6*^*/FM7c, sn*^*+*^ (BDSC Stock 2497); *brk*^*M68*^/*FM7c*-*ftz*-*lacZ* (Jazwinska et al., 1999). *y*^*1*^ *w*^*67c23*^ flies were used as controls throughout.

### CRISPR-Cas9 genome editing and PhiC31 reintegration

The *sog*^*attP*^ CRISPR *Drosophila* line was generated by CRISPR/Cas9 with homology directed repair (HDR) in a two-step CRISPR approach (Baena-Lopez et al., 2013; Hoppe & Ashe, 2021). For a detailed outline of the strategy see (Hoppe & Ashe, 2021). PAM sites (NGG) located either side of the *sog* start codon were identified using the CRISPR OptimalTarget Finder tool on the flyCRISPR website (Gratz et al., 2014). Two guide RNA sequences were designed 3 nucleotides upstream of the selected PAM sites to target these sites for Cas9 nuclease digestion and the creation of double stranded breaks. For oligonucleotides sequences encoding sense and antisense strands for guide sequence see Table S1. Complementary guide oligonucleotides were annealed and inserted into the pU6-Bbs1-gRNA plasmid (RRID:Addgene_45946, (Gratz et al., 2013)) as described (Hoppe & Ashe, 2021). Homology arm (HA) sequences were obtained from *Drosophila* genomic DNA (BL51324) by PCR. Homology arms were inserted into the pTV^Cherry^ donor plasmid (Drosophila Genomics Resource Center, DGRC_1338) (Baena-Lopez et al., 2013) at SpeI and KpnI restriction sites. Donor plasmids and gRNA plasmids were injected into Cas9 expressing embryos (BL51324) at the University of Cambridge Fly Injection Facility. Flies that developed from injected embryos were crossed according to (Hoppe & Ashe, 2021), using *y*^*1*^ *w*^*67c23*^ and *brk*^*M68*^/FM7c-*ftz*-*lacZ* (Jazwinska et al., 1999) stocks. The *white*+ marker was removed by crossing *sog*^*attP*^ females with males which carry FM7c on the X and Cre-recombinase on the third chromosome. The *sog*^*attP*^ CRISPR mutation results in a deletion of 800bp (ChrX:15,625,466, - 15,626,265, dm6), including the endogenous *sog* start codon and signal sequence, that is replaced by 103 nucleotides encoding attP and LoxP sites. Reintegration plasmids were generated from the RIV^White^ plasmid (gift from the Vincent lab; DGRC 1330). A partial *sog* 5’UTR (p5’UTR) sequence (source: pBS-Sog-CDS, (Ashe & Levine, 1999)), followed by the *sog* CDS, mNeonGreen the *sog* 3’UTR (source: pBS-Sog-CDS, (Ashe & Levine, 1999)), was inserted between the RIV^White^ attB and LoxP sites. To summarise the construction of these plasmids, the partial 5’UTR and the *sog* CDS were ligated together in a pAc5.1/V5-His (ThermoFisher, V411020) vector, as were the mNeonGreen and *sog* 3’UTR sequences, using In-fusion cloning (Takara Biosciences). A linker sequence was added downstream of the *sog* CDS. The p5’UTR-*sog* CDS-linker, and mNeonGreen-3’UTR sequences were inserted into RIV^White^ using In-Fusion cloning. To create the *sog*^*C27,28S*^ mutant, Cys 27 and 28 were replaced by two Ser residues with In-Fusion cloning. Reintegration vectors were injected with a PhiC31 encoding plasmid into embryos of the *sog*^*attP*^ CRISPR stock. Female flies that developed from the injected embryos were crossed to *y*^*1*^ *w*^*67c23*^ males and *w*+ offspring were crossed to *brk*^*m68*^/FM7c *ftz-lacZ* to balance. Successful PhiC31 mediated recombination was confirmed by sequencing genomic DNA. Flies in which successful PhiC31 recombination events had occurred were back-crossed to each other to make homozygous stocks. The *w*+ marker gene was not removed from the *sog-mNG* and *sog*^*C27,28S*^*-mNG* stocks generated here. This could be done by Cre-recombinase if necessary for future work. See Table S1 for primers and oligonucleotide sequences. SnapGene viewer software (from Insightful Science; available at snapgene.com) was used to scan DNA sequences for ORFs.

### Viability and lethality assays

For viability assays, 50 embryos were placed on an apple juice agar plate which was transferred into a food bottle. Embryos were incubated at 25°C in the bottle. The number of pupae and adults were counted. For the *sog* recue assay, virgins of *y*^*1*^ *sogS6/FM7c, sn+* (BDSC Stock 2497) or *sog*^*attP*^*/FM7* were crossed to male *yw, sog-mNG;;* or *sog*^*C27,28S*^;; flies. The number of FM7 and non-FM7 female offspring were counted to assess the degree of rescue from each genotype in the presence of the mutant *sog* allele.

### Wing dissection

Adult flies were incubated at 18°C or 25°C for 24 hours, before transfer to a fresh vial. Flies were then allowed to lay eggs before being discarded. Embryos were allowed to develop to adulthood at the designated temperature condition. Wings were then removed from adult females, placed on a slide and washed briefly in isopropanol. Wings were mounted in DPX mounting media (Fisher D/5319/05) under a No.1 coverslip. Samples were imaged with a light microscope (Zeiss Axioskop) with a 5X objective (Zeiss CP-Achromat 5X/0.12). Images were acquired with Agilent OpenLab 2.2.2 software. For analysis, Fisher’s exact tests were performed on raw count data in RStudio.

### *In situ* hybridisation and immunofluorescence

Embryos (2-4h) were collected and stained by RNA *in situ* hybridisation with *sog*-digoxygenin-UTP, *Race*-Biotin-UTP, *lacZ*-digoxygenin-UTP or mNeonGreen-biotin-UTP probes as described (Hoppe et al., 2020; Kosman et al., 2004). A mNeonGreen-biotin-UTP probe was synthesised as described (Kosman et al., 2004) with primers listed in Table S1. Antibodies used were mouse anti-biotin (1:250, Roche Cat# 1297597), sheep anti-digoxigenin Fab fragments antibody, AP conjugated (1:200, Roche Cat# 11093274910 RRID:AB514497), donkey anti-mouse IgG secondary antibody, Alexa Fluor 647 (1:500, ThermoFisher Cat# A-31571, RRID:AB162542), and donkey anti-sheep IgG secondary antibody, Alex Fluor 488 (1:500, ThermoFisher Cat#A-11015, RRID: AB_2534082). For pMad immunostaining, anti-Smad3 (phospho S423 + S425) [EP823Y] (1:500, Abcam Cat# ab52903, RRID: AB_882596) primary antibody and Donkey anti-rabbit IgG secondary antibody, Alexa Fluor 647 (1:500, ThermoFisher Cat#A-31573, RRID: AB_2536183) were used. To stain embryo nuclei, samples were incubated with DAPI (1:1000, NEB 4083). Samples were mounted in ProLong™ Diamond Antifade Mountant (ThermoFisher P36961).

### smFISH (Stellaris), smiFISH and immunofluorescence

For smFISH, 2-4h or 1-3h embryos were processed as described (Hoppe et al., 2020) with anti-*sog* Stellaris (Fig. 2C, Fig. S1), or *sog* single molecule inexpensive FISH (smiFISH) (Fig. S4) (Tsanov et al., 2016), *ush* Stellaris, *lacZ* Stellaris, and *Race* smiFISH probes (Tsanov et al., 2016). *ush* probe sequences are available from (Hoppe et al., 2020), while *sog* Stellaris, *sog* smiFISH, *lacZ* Stellaris, and *Race* smiFISH probe sequences are provided in Table S2. *Race* smiFISH probes were annealed to a 570-conjugated Y-FLAP (Tsanov et al., 2016). For immunostaining against mNeonGreen, mouse anti-mNeonGreen [32F6] (1:500, ChromoTek Cat# 32f6-100, RRID: AB_2827566), mouse anti-Spectrin (1:50, DSHB Cat# 3A9 (323 or M10-2), RRID: AB_528473) and donkey anti-mouse IgG secondary antibody, Alexa Fluor 488 (1:1000, Thermo Fisher Scientific Cat# A-21202, RRID: AB_141607) primary and secondary antibodies were used. To stain nuclei, samples were incubated with DAPI (1:1000, NEB 4083). Samples were mounted in ProLong™ Diamond Antifade Mountant (ThermoFisher P36961).

### Imaging stained embryos

Fixed embryos stained with *Race* and *lacZ* RNA *in situ* hybridisation probes were imaged with a Leica TCS SP5 AOBS inverted microscope using a HCX PL APO lambda blue 20.0×0.70 IMM UV oil objective. The following confocal settings were used: pinhole 1 airy unit, scan speed 400Hz unidirectional, 512×512 pixel format, Z step size of 1.5 μm at 8 bit and 1.3 zoom. Images shown are maximum intensity projections. Images were deconvolved with Huygens Professional software (SVI, Scientific Volume Imaging, RRID:SCR 014237).

*sog* Stellaris early nc14 embryos, co-stained with *lacZ* stellaris and anti-Spectrin antibody, were imaged in a Leica TCS SP8 AOBS confocal microscope with a HC PL APO CS2 40x/1.30 oil objective with 0.75 zoom to screen embryos that did not stain with *lacZ* smFISH. The settings used were as follows, pinhole 1 airy unit, scan speed 400Hz bidirectional, format 2048×512 pixels, at 8 bit. Images were collected using hybrid detectors using the white light laser with 647 nm (20%), 548 nm (20%), 405 nm (6%). To age embryos using the anti-Spectrin antibody stain, at the centre of the embryo stacks of 5-10 images (Z step size = 0.3 μm) were collected with hybrid detectors using the white light laser with 548 nm (20%), 488 nm (12%) 405 nm (6%). To collect images for analysis embryos were imaged with a HCX PL Apo 63x/1.40 oil objective and 0.75 X zoom. The confocal settings used were as follows, pinhole 1 airy unit, scan speed 400Hz bidirectional, format 2048×512 pixels, at 8 bit and Z step size 0.3 μm. Images were collected using hybrid detectors using the white light laser with 548 nm (20%), 405 nm (6%) with 3X line accumulation. Raw images were deconvolved with Huygens Remote Manager software v3.7.1 (SVI). Images shown are maximum intensity projections.

*sog* smiFISH stage 6 embryos were imaged on a Leica TCS SP8 AOBS confocal microscope with a HCX PL Apo 63x/1.40 oil objective with 1 x zoom. The confocal settings used were as follows, pinhole 1 airy unit, scan speed 400Hz bidirectional, format 2048×2048 pixels, at 8 bit and Z step size 0.3 μm. Images were collected using hybrid detectors using the white light laser with 548 nm (20%), 405 nm (9.8%) with 3 X line accumulation. Raw images were deconvolved with Huygens Remote Manager software v3.7.1 (SVI). Images shown are maximum intensity projections. Raw images were deconvolved with Huygens Remote Manager software v3.7.1 (SVI). Samples subject to FISH with *sog*-digoxygenin and mNeonGreen-biotin RNA probes were imaged on a Leica TCS SP8 AOBS confocal microscope with a HCX PL Apo 63x/1.40 oil objective and 0.75 zoom. The confocal settings used were as follows, pinhole 1 airy unit, scan speed 400Hz unidirectional, format 2048×700 pixels, at 8 bit, and Z step size 0.35 μm. Images were collected using hybrid detectors using the white light laser with 647 nm (14%), 488 nm (14%), 405 nm (14%), and 4X line averaging. Images were deconvolved with Huygens Professional software. Cropped squares from the 2048×700 images are presented and the images shown are single Z slices.

*sog* smFISH and anti-mNeonGreen immunofluorescence samples were imaged on a Leica TCS SP8 AOBS confocal microscope with a HC PL APO CS2 40x/1.30 oil objective and 0.75 zoom. The confocal settings used were as follows, pinhole 1 airy unit, scan speed 400Hz, format 2048×2048 pixels, at 8 bit and 0.35 μm Z step size. Images were collected using hybrid detectors using the white light laser with 647 nm (22%), 548 nm (23%), 488 nm (20%), 405 nm (14%), and 6X line averaging. Images were deconvolved with Huygens Professional software. Images shown are maximum intensity projections (∼3.5 μm depth).

Embryos stained for pMad were imaged on a Leica TCS SP8 AOBS confocal microscope with a HCPL APO CS2 40x/1.30 oil objective and 0.75 zoom. The confocal settings were as follows; pinhole of 1 airy unit, unidirectional scanning at 400Hz, 1024×1024 pixel format at 8 bit and Z step size of 0.35 μm. Images for these samples were collected with hybrid detectors using a white light laser with 488nm (20%), 647 (27%), 405nm (12%) and 4X line averaging. Raw images were deconvolved with Huygens Remote Manager software v3.7.1 (SVI). Images for *Race* FISH co-stain are not shown.

*ush* smFISH and *Race* smiFISH samples were imaged on a Leica TCS SP8 AOBS confocal microscope with a HCX PL Apo 63x/1.40 oil objective and 1X zoom. The confocal settings used were as follows, pinhole 1 airy unit, scan speed 400Hz bidirectional, format 3144×3144 pixels, at 8 bit and Z step size 0.3 μm. Images were collected using hybrid detectors using the white light laser with 647 nm (22%), 548 nm (22%), 405 nm (6.1%) with 3X line accumulation. Raw images were deconvolved with Huygens Remote Manager software v3.7.1 (SVI). Images shown are maximum intensity projections.

### smFISH/smiFISH image analysis

To age *sog*^*attP*^ and *y*^*1*^ *w*^*67c23*^ control embryos stained with anti-*sog* and anti-*lacZ* smFISH and anti-Spectrin antibody, maximum intensity projections of images were made, and the length of the cell membrane ingression (Spectrin antibody stain) was measured (Calvo et al., 2021). Embryos with cell membranes of 3.5-5.5 μm were used for analysis. Only male embryos were analysed. *y*^*1*^ *w*^*67c23*^ males were identified by the number of *sog* transcription sites, and *sog*^*attP*^ males were identified by the absence of *lacZ* smFISH stain.

Quantitative analysis of *ush* and *Race*, and *sog* smFISH/smiFISH images was performed in Imaris 9.2 (Bitplane, Oxford Instruments). For efficiency, analysis of *ush*/*Race* images was performed for only the central 1048 × 3144 area of the 3144×3144 images. For *sog* smFISH data, the entire 2048×512 images were analysed. For quantification of *ush, Race*, or *sog* mRNA number, individual mRNAs were detected with the ‘spots’ function. Spots of diameter 0.3 μm (X/Y direction) and 0.8 μm (Z direction) were used. Nuclear locations were determined using the ‘surfaces’ function to identify nuclei based on DAPI staining. To determine the number of spots per cell, spots were assigned to surfaces using the spotMe_V2.py python script (Vinter et al., 2021) (script available at https://github.com/TMinchington/sass). Output from this analysis was processed in Rstudio to remove duplicated nuclei values and ‘NAs’. Data for the number of mRNAs for each of the three embryos analysed was pooled, and divided into 5 μm bins (approximate cell size) to permit calculation of the mean number of mRNAs at a given distance from the dorsal/expression domain midline. To account for imaging of embryos that were not perfectly lateral, data embryos stained with anti-*sog* smFISH, were cropped -60 and 80 μm from the expression domain midline, as these were the boundaries shared by all embryos. Binned data were plotted in Rstudio, while data for individual embryos were plotted in GraphPad Prism 9 (RRID:SCR 002798).

For analysis of *sog* expression in *sog-mNG* and *sogC27,28S-mNG* and control embryos, *sog* smiFISH images were analysed in Fiji ImageJ (Schindelin et al., 2012). To quantify the height of the *sog* expression domain, maximum intensity projections of the imaging stacks were made, and the maximum vertical number of *sog* positive cells was counted in a 25 μm region of interest (ROI) located 50 μm to the posterior of the cephalic furrow. To quantify *sog* expression level within *sog* positive cells in sum of slices projected images, a 40 × 40 μm ROI was drawn 50 μm away from the cephalic furrow and bordering the ventral edge of the *sog* expression domain. The mean grey value of the ROI was calculated. To correct for background fluorescence, the mean grey value along a 20 μm line situated outside the sog expression domain was calculated. This value was subtracted from the mean grey value within the ROI. Statistical analysis was performed in GraphPad Prism 9 (RRID:SCR 002798).

### pMad stain analysis

To quantify pMad distribution in stage 6 embryos, the width of the pMad stripe was measured as the number of pMad positive nuclei in maximum intensity projected images. Analysis was performed in Fiji ImageJ (Schindelin et al., 2012). The data were plotted and the statistical test was performed in GraphPad Prism (RRID:SCR 002798).

### Amnioserosa counts

Fixed embryos were stained with mouse anti-Hnt 1G9 (1:40, DSHB Cat# 1g9, RRID: AB_528278) and Anti-Mouse IgG (H+L) AP Conjugate S372B 1:500 (Promega) by standard techniques (Kosman et al., 2004). Stage 11 embryos were imaged on a Leica DM600B with a 20x objective using brightfield. Total amnioserosa cells were counted on Image J using the Cell Counter plugin. 50 embryos across 3 biological repeats were analysed per genotype. Counts were plotted in GraphPad Prism 9 (RRID:SCR 002798).

## Acknowledgements

We thank Drs Michael Lockhart-Cairns and Alan Godwin for helpful discussions and support with EM data collection and analysis, Dr Thomas Jowitt (University of Manchester Biomolecular Analysis Facility) for AUC data collection and analysis, the Bloomington Drosophila Stock Centre for flies, JP Vincent lab for plasmids, and the Cambridge Fly Facility for microinjections. We are grateful to the Baron lab for wing imaging training and Osamu Shimmi for helpful discussions. We also thank staff of The University of Manchester Bioimaging Facility, and Fly Facility Manchester for their support and staff of the FBMH EM Core Facility (RRID:SCR_021147) for their assistance.

## Author Contributions

Conceptualisation, S.L.F, C.B, and H.L.A.; Investigation, S.L.F, C.S.; Writing, S.L.F. and H.L.A.; Editing, S.L.F, C.S., C.B, and H.L.A.

## Competing Interests

None.

## Funding

This research was supported by a BBSRC project grant (BB/V008099/1) to H.A. and C.B and a BBSRC DTP PhD studentship (BB/M011208/1) to S.F.

## Notes

### Competing Interest Statement

The authors have declared no competing interest.

