## Supplemental data for "Modelling the structure of Short Gastrulation and generation of a toolkit for studying its function in *Drosophila*"

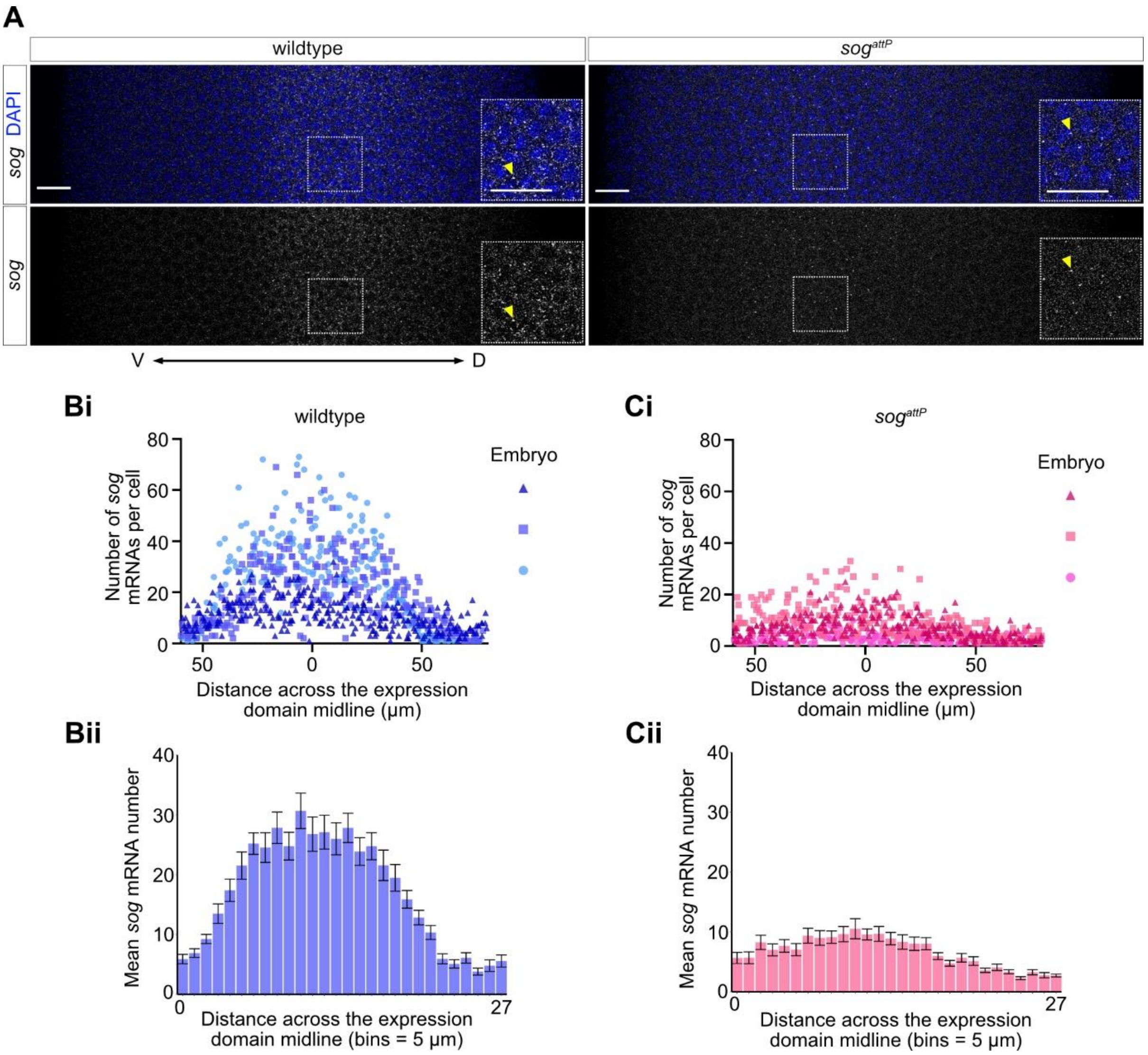


**Figure S1. *sog* is transcribed at low levels in early nc14 *sog*$^{attP}$ embryos.**

A) Confocal images (maximum intensity projections) showing *sog* mRNAs in early nc14 wildtype and *sog*$^{attP}$ male embryos stained with *sog* smFISH probes and the DAPI nuclear marker. A merged image and single smFISH channel are shown for clarity. Spectrin antibody staining was used to measure membrane ingression to age the embryos (data not shown). Scale bar = 15 µm. Insets are enlarged sections of the *sog* expression domain (box), scale bar = 15 µm). Yellow arrowheads indicate transcription sites.

Bi) The numbers of *sog* transcripts per cell in three wildtype embryos were quantified and are plotted against distance from the *sog* expression domain midline (µm, - 60 = ventral edge). Data for each embryo is presented on the same graph in a shade of blue with a triangle, square, or circle symbol.

Bii) Data for each wildtype embryo were combined and binned according to distance across the *sog* expression domain (bins = 5 µm, each bin approximately corresponds to a nuclear width, bin 0 = ventral edge). Mean *sog* mRNA number per bin is plotted against bin number. Error bars = mean ± SEM.

Ci) As for Bi, but for *sog* mRNA counts in three *sog*$^{attP}$ embryos. Each embryo is represented by a shade of pink and either a triangle, square or circle symbol.

Cii) As for Bii, but for *sog*$^{attP}$ embryos.

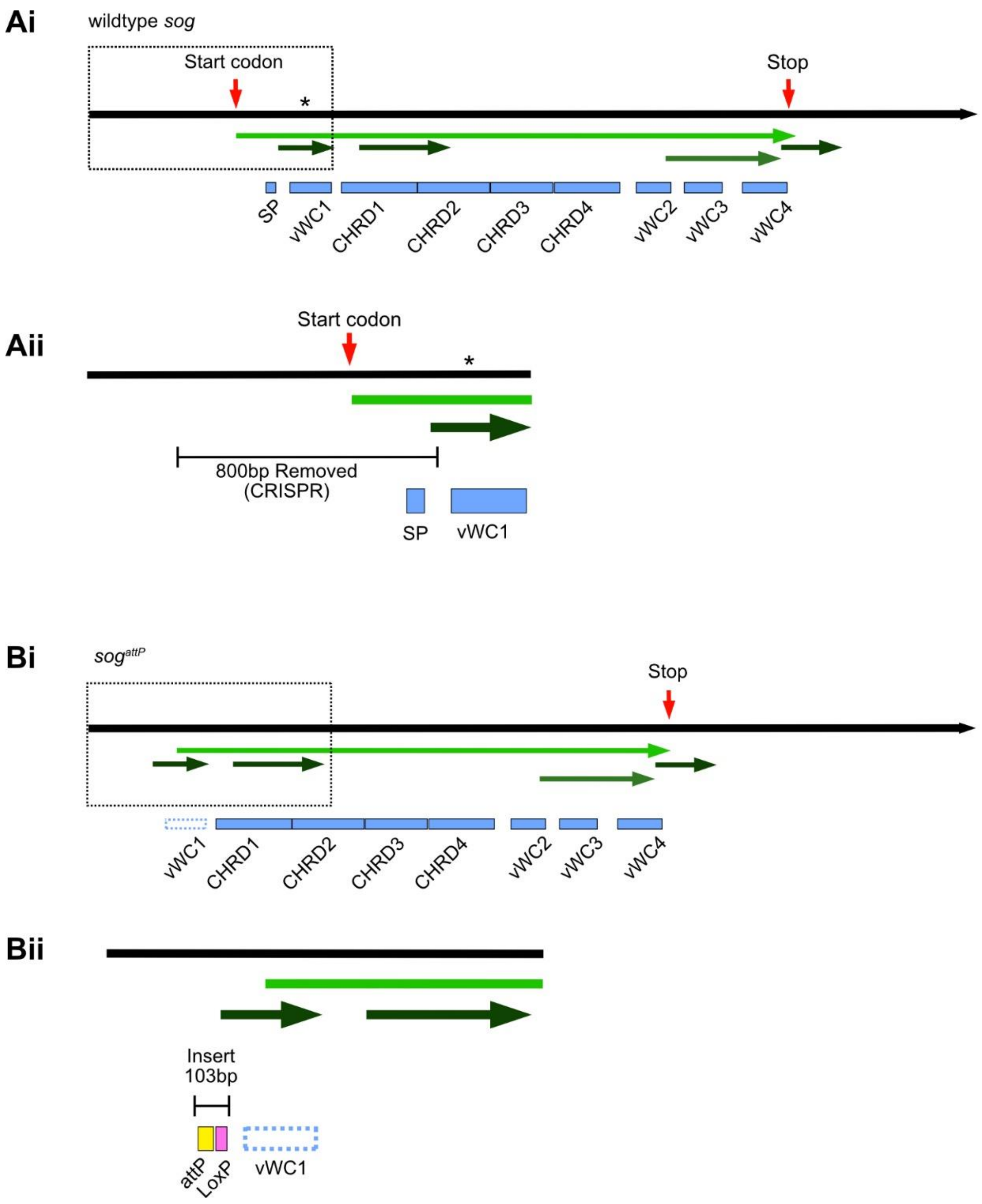


**Figure S2. Potential ORFs in the wild type *sog* and CRISPR modified *sog*$^{attP}$ transcripts.** Ai) Schematic showing the wildtype *sog* transcript (isoform E, (Larkin et al., 2021)). ORFs are shown by green arrows, with the shade of green indicative of the reading frame. A red arrow indicates the endogenous start and stop codons of the *sog* coding sequence. The asterisk marks the position where a truncated Sog ORF initiates following the CRISPR deletion (see Bi).

Aii) An enlarged view of the outlined region in Ai) showing the 800 bp sequence removed by CRISPR-Cas9 and the locations of the predicted signal peptide (SP) and vWC1 domain (blue boxes).

Bi) Schematic showing the locations of predicted ORFs (green arrows) in the predicted $sog^{attP}$ transcript (*sog* isoform E).

Bii) An enlarged view of the outlined region in Bi) showing the location of the 103 bp insert, containing attP and LoxP sequences (yellow and pink boxes, respectively), introduced into the sequence by CRISPR-Cas9 mediated homology directed repair, and the vWC1 domain. Due the CRISPR-Cas9 genome editing, none of the predicted ORFs contain the Sog SP, and therefore any translated protein would not enter the secretory pathway.

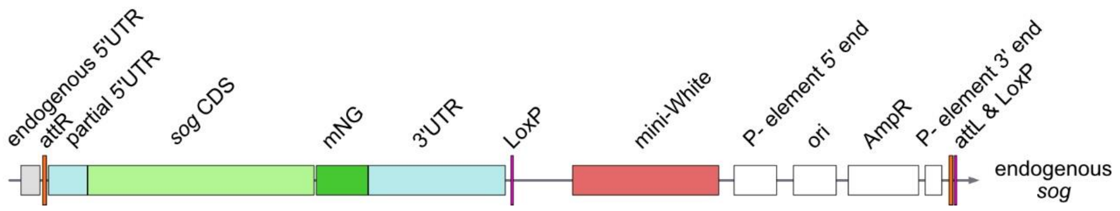


**Figure S3. The *sog* locus after CRISPR-Cas9 editing and reintegration of *sog-mNG* sequences.**

The *sog* CDS and entire RIV white plasmid (Baena-Lopez et al., 2013) were inserted into the genome by phiC31 recombination, resulting in insertion of a 12,565 bp sequence. Due to the location of the CRISPR-Cas9 cut sites, some of the *sog* 5'UTR was removed in the initial CRISPR event. The grey box in the schematic represents the remaining endogenous 5'UTR, while the light blue box labelled 'partial 5'UTR' represents the 5'UTR sequence used to replace the 5'UTR sequences which were initially removed. The LoxP sequences (pink) located after the *sog* 3'UTR and P-element 3' end sequences permit removal of 6,081 bp after the 3'UTR by Cre-Lox recombination. However, this was not performed for this study.

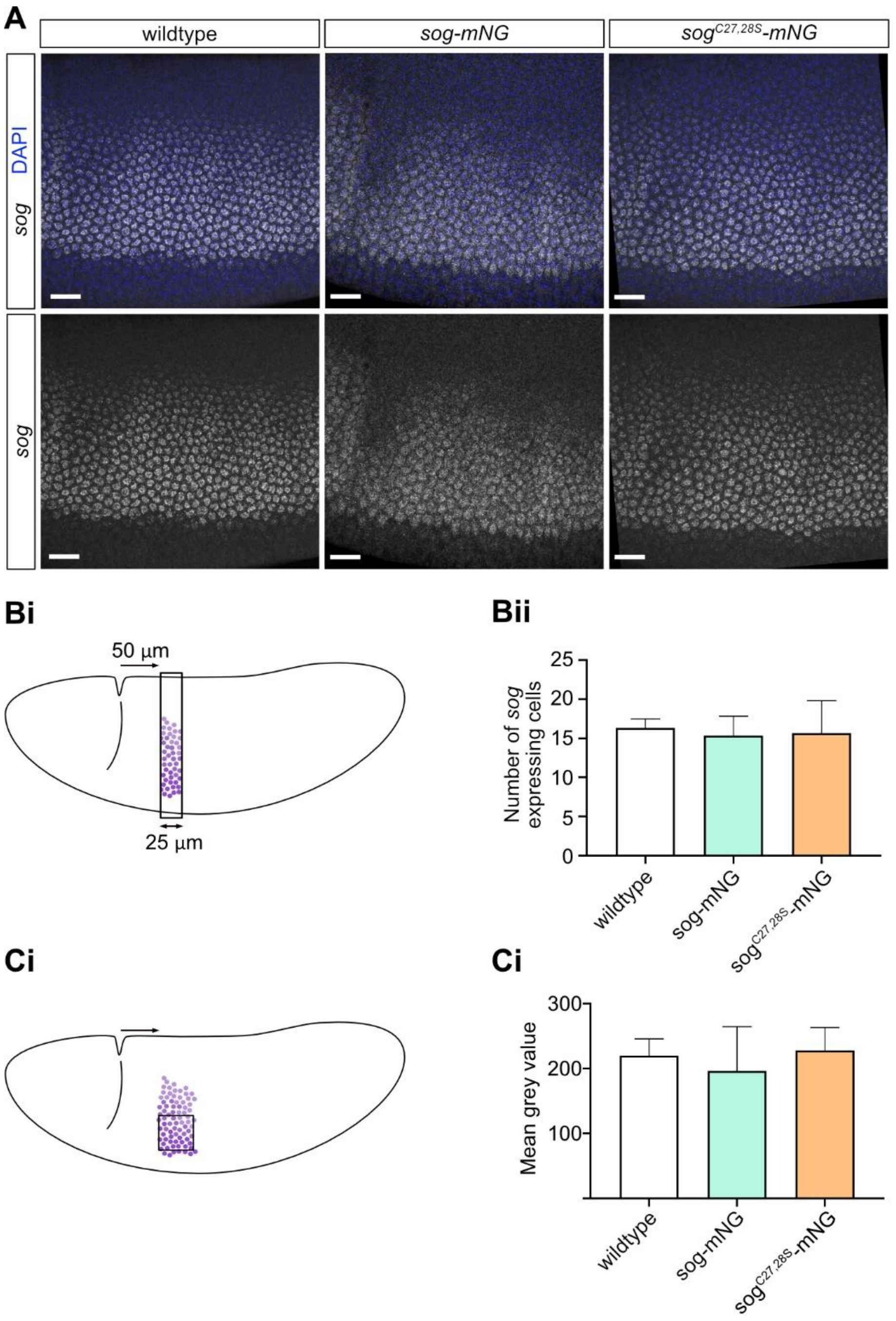


**Figure S4. *sog* expression in nc14 embryos is similar between genotypes.**

A) *sog* mRNA was detected in wildtype, *sog-mNG*, and *sog*$^{C27,28S}$*-mNG* early stage 6 embryos by smiFISH. Scale bar = 20 µm.

Bi) Cartoon showing the position of a region of interest (ROI) that was used to measure the height of the expression domain based on the maximum number of *sog* expressing cells. The ROI is located 50 µm posterior of the cephalic furrow.

Bii) Graph shows the maximum number of *sog* expressing cells in the ROI depicted in Bi. No significant difference in the height of the *sog* expression domain was found (One-Way ANOVA with Tukey’s multiple comparisons test, P>0.9). n = 3 embryos for each genotype, error bars show standard deviation.

Ci) Cartoon showing a 40x40 μm ROI, 50 μm to the posterior of the cephalic furrow, that was used to estimate *sog* expression level in early stage 6 embryos. Mean fluorescence intensity (based on grey value), of the ROI was measured, as clustering of *sog* mRNAs at this stage prevents the counting of absolute mRNA numbers as is possible in early nc14 embryos (see Fig S1).

Cii) Background subtracted fluorescence intensity (grey) values of the ROI are shown. No significant difference was detected between genotypes (One-Way ANOVA, Tukey’s multiple comparisons test, P>0.7). n = 3 embryos for each genotype, error bars show standard deviation.

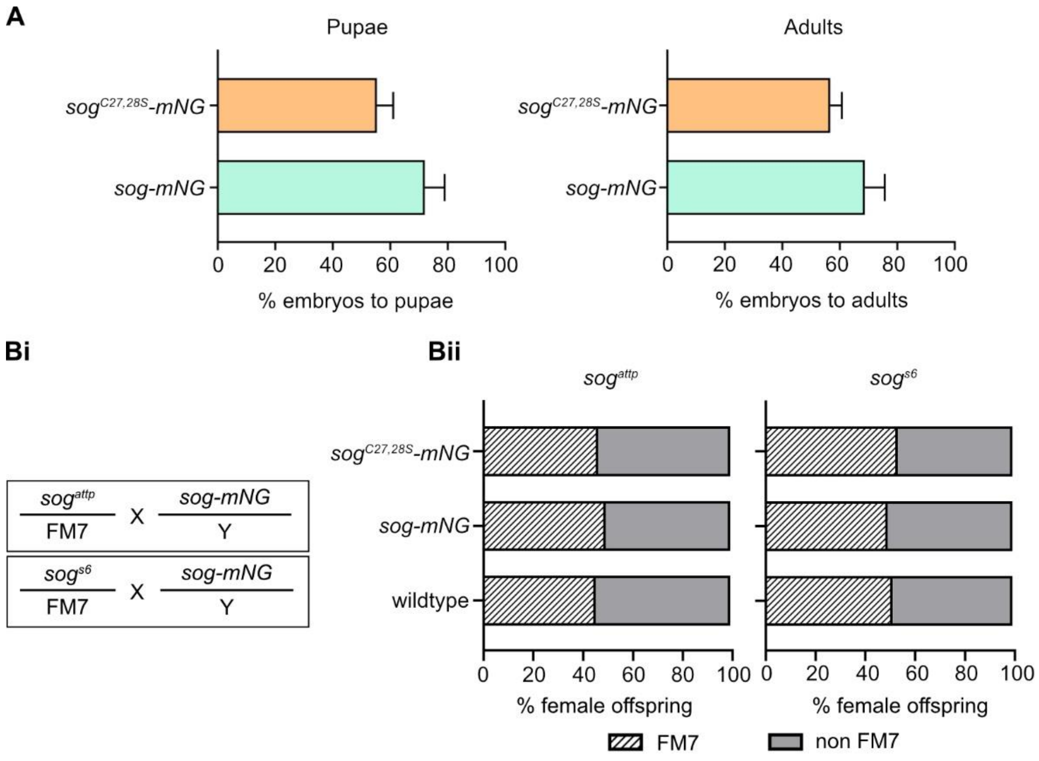


**Figure S5. Assessing viability of *sog-mNG* and *sog*$^{C27,28S}$*-mNG* flies.**

A) Graphs show the percentage survival of *sog-mNG* and *sog*$^{C28,28S}$*-mNG* embryos to pupal and adult stages. No significant difference was found between the proportion of *sog-mNG* and *sog*$^{C27,28S}$*-mNG* flies that survived to pupae (p=0.14) and adulthood (p=0.21) (t-test, unpaired, two-tailed. t-tests were performed on raw data).

Bi) Overview of the crosses used to compare the ability of the *sog-mNG* and *sog*$^{C27,28S}$*-mNG* alleles to rescue either the strong *sog*$^{S6}$ loss of function or *sog*$^{attP}$ null alleles. Wildtype males were also tested as a control, only the *sog-mNG* allele is shown in the crossing scheme for simplicity. The number of female FM7 and non-FM7 progeny were counted. For female progeny that do not carry FM7, the *sog* allele has successfully rescued the *sog*$^{attP}$ or *sog*$^{s6}$ loss of function alleles.

Bii) The percentage of FM7 (striped bars) and non-FM7 (grey bars) progeny from the crosses described in Bi are shown. There is no significant association between the number of FM7/non-FM7 offspring and the genotype of fathers for crosses with *sog*$^{attP}$ ($X^2$ (df = 2, N = 1764) = 2.07, p = 0.36) and *sog*$^{S6}$ ($X^2$ (df = 2, N = 1596) = 1.71, p = 0.43) mothers. Statistical test was performed on raw data. Percentage frequency of the phenotypes is shown in the figure for simplicity.

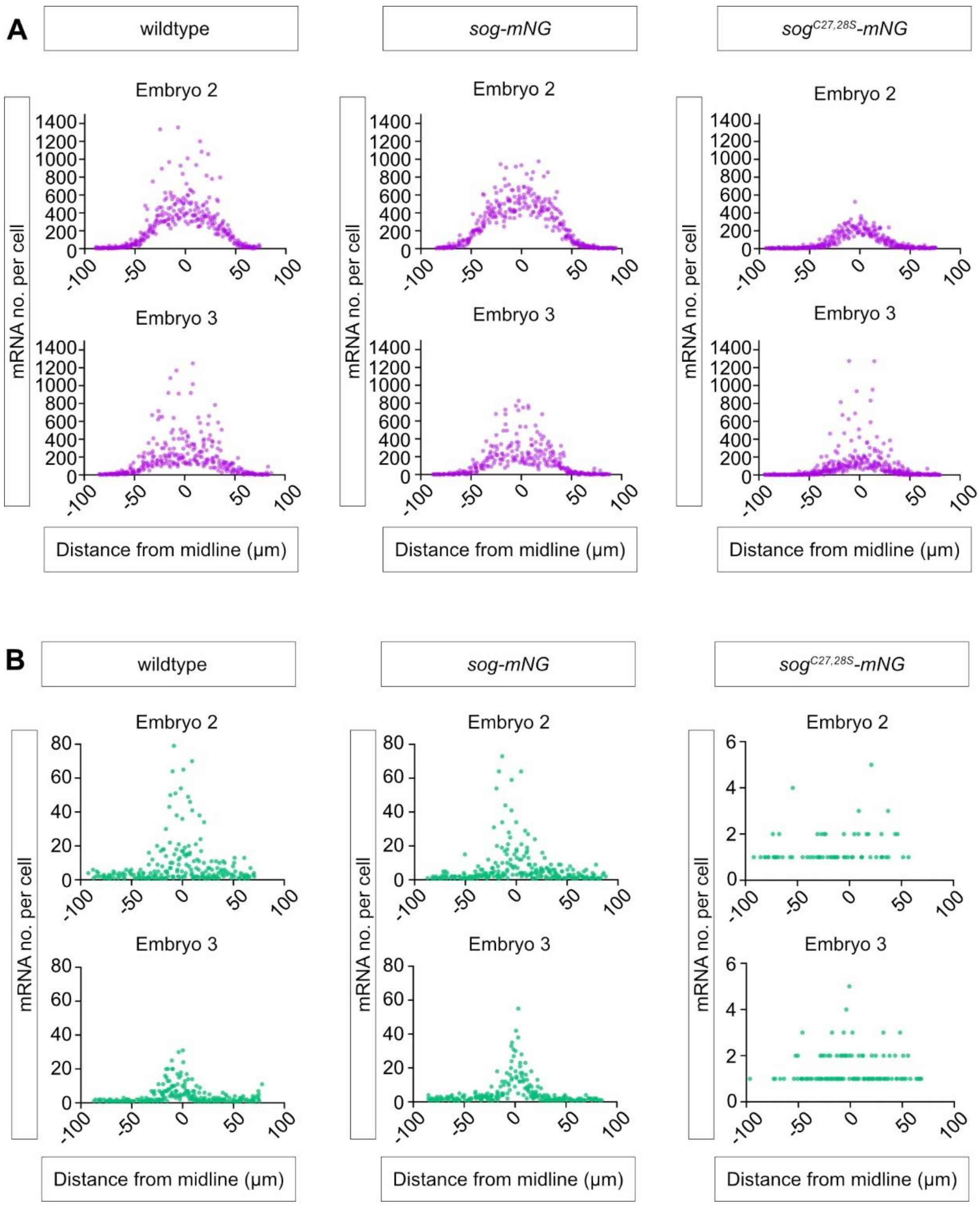


**Figure S6. Quantification of *ush* and *Race* expression.**

(A) The numbers of *ush* transcripts per cell plotted against distance from the dorsal midline (µm, 0 = midline) for the additional biological repeats relating to Fig.4.

(B) As in (A), but for *Race* mRNAs.

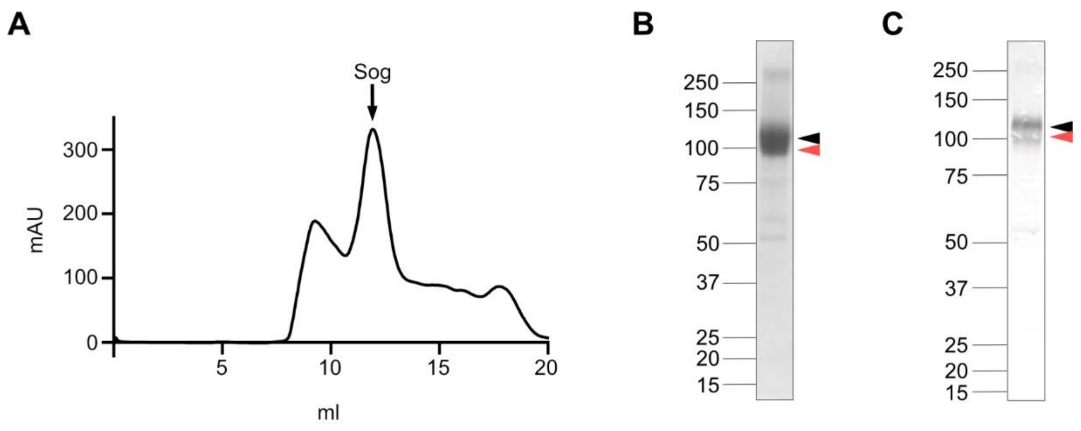


**Fig. S7. Purification of Sog by size exclusion chromatography.**

A) Following affinity chromatography using a His-tag, Sog was purified by SEC. The peak Sog-containing fraction, indicated by an arrow, was subjected to a second round of SEC (main text, Fig.6) prior to electron microscopy.

B) Reduced SDS-PAGE gel showing Sog from the peak fraction in A. The black arrow indicates full-length Sog and the red arrow indicates a Sog cleavage product.

C) Anti-His western blot of the same sample shown in B. Arrows are as in B.

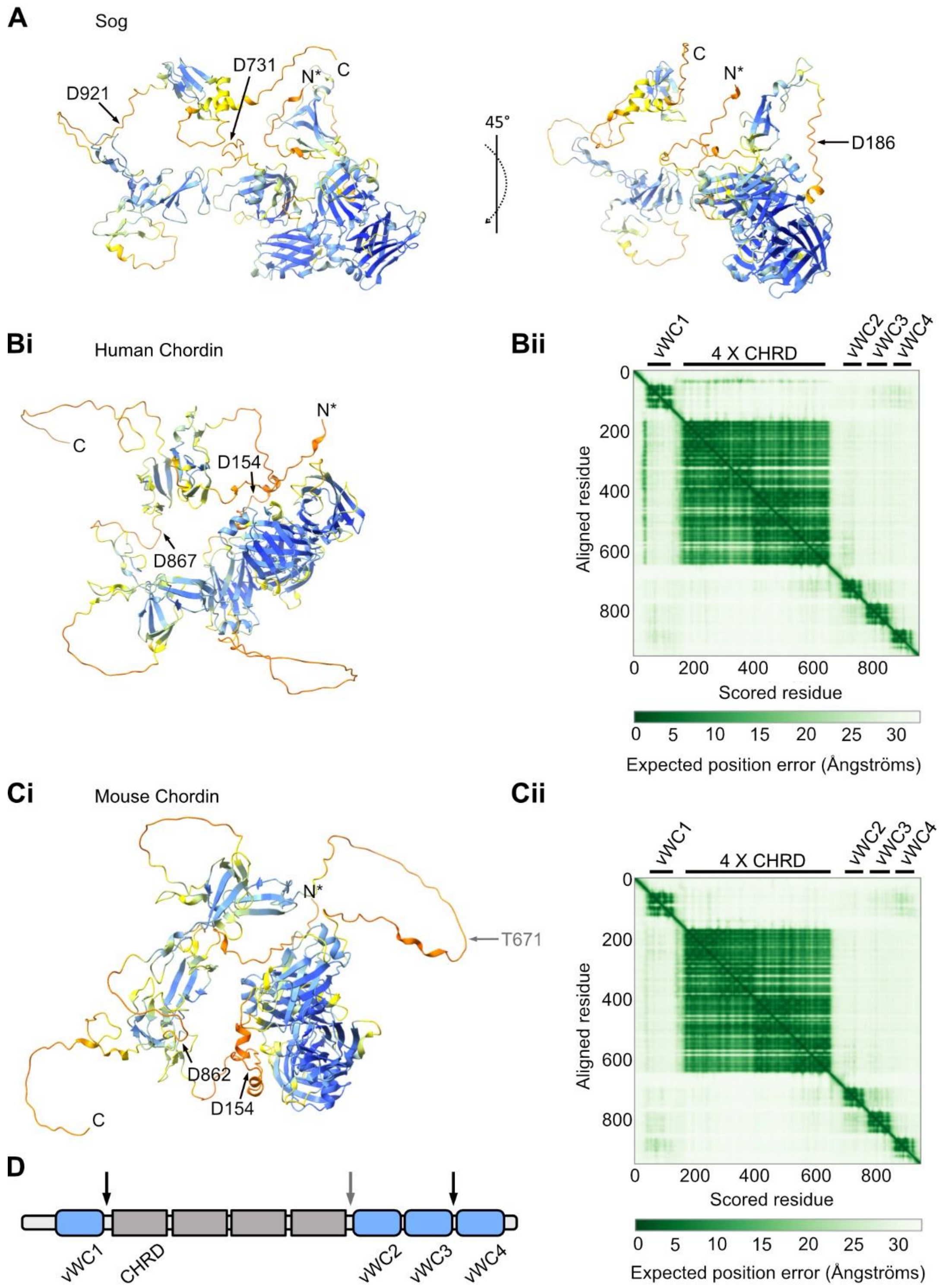


**Fig. S8. Interdomain regions that contain the Sog/Chordin Tolloid cleavage sites are flexible.**

A) AlphaFold2 structure prediction for Sog (Jumper et al., 2021; Varadi et al., 2022), colour coded according to per residue confidence (pLDDT) score (dark blue, very high confidence; light blue, confident; yellow, low confidence; orange, very low confidence). Tld cleaves Sog before D921, D731, and D186 (arrows) (Peluso et al., 2011; Shimmi & O’Connor, 2003) . In the figure shown, the Sog N-terminus is truncated at R79 (N*).

Bi) AlphaFold2 structure prediction for human Chordin (UniProt Q9H2X0), coloured as for Sog in (A). BMP1/Tld-like protease cleavage sites are annotated (arrows) (Scott et al., 1999). The predicted Chordin structure has been truncated following the signal peptide at G26 (N*).

Bii) Predicted aligned error (PAE) plot for the AlphaFold human Chordin prediction. Approximate boundaries for vWC and CHRD (4 X CHRD) domains are indicated by black bars above the heatmap.

Ci) AlphaFold2 structure prediction for mouse Chordin (Uniprot Q9Z0E2), coloured as for human Chordin and Sog. Tld cleavage sites are indicated by arrows. The predicted structure has been truncated after the signal peptide at G26 (N*). Two cleavage sites (D154 and D862) are conserved in human Chordin. A third Tld cleavage site (T671, grey arrow) in mouse Chordin is also used in the presence of Tsg *in vitro* (Scott et al., 2001).

Cii) Predicted aligned error (PAE) plot for the AlphaFold mouse Chordin prediction.

D) Cartoon of Chordin (not to scale), showing the vWC and CHRD domains in relation to Tld cleavage sites present in both mouse and human Chordin (black arrows), and the cryptic Tld target site (grey arrow) found in mouse Chordin in the presence of Tsg.

**Table. S1. Primer and oligonucleotide sequences.**

| Plasmid | Primer | Sequence (5'-3') |
|---|---|---|
| pU6-BbsI-chiRNA-guides | Guide 1 Forwards | [Phos]CTTCGAGTCGATCTCGTATGAGGA |
| | Guide 1 Reverse | [Phos]AAACTCCTCATACGAGATCGACTC |
| | Guide 2 Forwards | [Phos]CTTCGCATGCGCCGCTCATGTTCG |
| | Guide 2 Reverse | [Phos]AAACCGAACATGAGCGGCGCATGC |
| pTV-Cherry-homology-arms | Homolog arm 1 Forwards | CTAGCACATATGCAGGTACCTTTAAGATTGTCAGCATTGCA |
| | Homolog arm 1 Reverse | AGTTGGGGCACTACGGTACCATTACGACAACGCGACTTTT |
| | Homology arm 2 Forwards | CGAAGTTATCACTAGTAGTCCGACACGGGCAGGC |
| | Homology arm 2 Reverse | GGAGATCTTTACTAGTGCAACTCGGGAACATAATAG |
| pAC-p5UTR-Sog CDS-link | KpnI p5UTR F | GGTACCTCATACGAGATCGACTCTATTTTCC |
| | NotI SogCDS R | GCGGCCGCGCTGGAGGATCGCTGCT |
| | Linker sense | [Phos}GGCCGCCGGTGGTGGTGGAAGTGGCGGAGGTGGAGGGCC |
| | Linker antisense | [Phos]CTCCACCTCCGCCACTTCCACCACCACCGGC |
| pAC-mNeonGreen-sog3UTR | Kpn1 NG F | GGTACCATGGTGAGCAAGGGCG |
| | NotI NG R | GCGGCCGCTTACTTGTACAGCTCGTCCATG |
| | pACng3UTRinfFwd | CAAGTAAGCGGCCGCGCGGCTCCACGTGACGGAT |
| | pAC3UTRinfRev0818 | ACCTTCGAAGGGCCCATGGGTATATTTCGAATATATTTTGTCTATATTTC |
| pAC-p5UTR-sogCDS-link-mNeonGreen-3UTR | ApaI NG F | GGGCCCATGGTGAGCAAGGGCG |
| | BstBI 3UTR R | TTCGAAATGGGTATATTTCGAATATATTTTGTCTATATTTCAATTTA |
| RIV-p5UTR-sog-link- | RIV p5UTR inffwd 250418 | GGCGCGTACTCCACGAATTCTCATACGAGATCGACTCTAT |
| | Linker rev20818 | TCCACCTCCGCCACT |

| | | |
|---|---|---|
| mNeonGreen-3UTR | Link-NGinfFwd0818 | GTGGAAGTGGCGGAGGTGGAATGGTGAGCAAGGGC |
| | RIV 3UTR infrev 250418 | GCGGCCGCTCCGGAGAATTCATGGGTATATTTCGAATATA |
| RIV-p5UTR-sogPALM-link-mNeonGreen-3UTR | RIV p5UTR inffwd 250418 | GGCGCGTACTCCACGAATTCTCATACGAGATC GACTCTAT |
| | palm mut rev 161018 | GCGGCGTCCTCGCTGTGAGAAGAGCTCCTTTCCAGGAGC |
| | palm mutfwd3 1218 | TCTCACAGCGAGGACGCCGC |
| | RIV 3UTR infrev 250418 | GCGGCCGCTCCGGAGAATTCATGGGTATATTTCGAATATA |
| **Probe** | **Primer** | **Sequence (5'-3')** |
| mNeonGreen-biotin-UTP | T3 promoter, forward. | ATTAACCCTCACTAAAGGGAATGGTGAGCAAGGGCGAGGAG GAT |
| | T7 promoter, reverse | GAATTAATACGACTCACTATAGGGATTACTTGTACAGCTCG TCCATGCC |

**Table. S2. Sequences of the *Race* smiFISH, *lacZ* Stellaris, and *sog* Stellaris and exonic smiFISH probes. *Race* smiFISH probes fused to Quasar 570-conjugated Y-FLAP, sog exonic smiFISH probes fused to Quasar 570-conjugated Z-FLAP, and *sog*, *ush* (Quasar 570 conjugated) and *lacZ* (Quasar 670 conjugated) Stellaris probes were used for smFISH.**

| Probe name | No. | Probe 5'-3' |
|---|---|---|
| race Y FLAP | 1 | TTACACTCGGACCTCGTCGACATGCATTATCCGTTAATTGCCCTAA |
| race Y FLAP | 2 | TTACACTCGGACCTCGTCGACATGCATTCCCCAATTAAAGGCTACT |
| race Y FLAP | 3 | TTACACTCGGACCTCGTCGACATGCATTTCGCGAAACAGTTCGCCA |
| race Y FLAP | 4 | TTACACTCGGACCTCGTCGACATGCATTTCCGTGGTGTGTACTACA |
| race Y FLAP | 5 | TTACACTCGGACCTCGTCGACATGCATTCGACCATGTCCACCGAAA |
| race Y FLAP | 6 | TTACACTCGGACCTCGTCGACATGCATTACCACCGCAGAAGTGTTT |
| race Y FLAP | 7 | TTACACTCGGACCTCGTCGACATGCATTTCGCTTTTGGTACACAGT |
| race Y FLAP | 8 | TTACACTCGGACCTCGTCGACATGCATTCGATCGTAAGTGCAACCT |
| race Y FLAP | 9 | TTACACTCGGACCTCGTCGACATGCATTTGGTTGGTTCGATCTGTT |
| race Y FLAP | 10 | TTACACTCGGACCTCGTCGACATGCATTACTCTGGAAGTCACTTCA |
| race Y FLAP | 11 | TTACACTCGGACCTCGTCGACATGCATTCTCAAGTGATTGCCCACA |

| | | |
|---|---|---|
| race Y FLAP | 12 | TTACACTCGGACCTCGTCGACATGCATTGGATGACTCTGGGGTCAG |
| race Y FLAP | 13 | TTACACTCGGACCTCGTCGACATGCATTGGGCTAGCAGAAACAGTC |
| race Y FLAP | 14 | TTACACTCGGACCTCGTCGACATGCATTTACCGCCAAAGTAGCCAG |
| race Y FLAP | 15 | TTACACTCGGACCTCGTCGACATGCATTCTCCTTGACCAGCGCTTG |
| race Y FLAP | 16 | TTACACTCGGACCTCGTCGACATGCATTTACTCCTTGGCCTGTATC |
| race Y FLAP | 17 | TTACACTCGGACCTCGTCGACATGCATTCCTTGTTCAGATTCTCCA |
| race Y FLAP | 18 | TTACACTCGGACCTCGTCGACATGCATTGTTGGTTCGCTTGGCCAG |
| race Y FLAP | 19 | TTACACTCGGACCTCGTCGACATGCATTCAGGCAGCTTCGGTTTCC |
| race Y FLAP | 20 | TTACACTCGGACCTCGTCGACATGCATTTGATGTTGGAGCCATAGG |
| race Y FLAP | 21 | TTACACTCGGACCTCGTCGACATGCATTTTTCTTTTCGTTCTCGTC |
| race Y FLAP | 22 | TTACACTCGGACCTCGTCGACATGCATTCTCGGCGGATATCTCATT |
| race Y FLAP | 23 | TTACACTCGGACCTCGTCGACATGCATTTCCTTCATGAACTTGGCC |
| race Y FLAP | 24 | TTACACTCGGACCTCGTCGACATGCATTCTTGGTGGTATCACTGGC |
| race Y FLAP | 25 | TTACACTCGGACCTCGTCGACATGCATTATTGGTACGAGCGCCATT |
| race Y FLAP | 26 | TTACACTCGGACCTCGTCGACATGCATTAACTGGCGCTTGAGATCC |
| race Y FLAP | 27 | TTACACTCGGACCTCGTCGACATGCATTAGCCCAGTTTGGTTAGAG |
| race Y FLAP | 28 | TTACACTCGGACCTCGTCGACATGCATTGTCTTCAGGTAGAGCAGC |
| race Y FLAP | 29 | TTACACTCGGACCTCGTCGACATGCATTTCCAGCAGTTCGGCATAG |
| race Y FLAP | 30 | TTACACTCGGACCTCGTCGACATGCATTACTCCATGGCGGAGAGTG |
| race Y FLAP | 31 | TTACACTCGGACCTCGTCGACATGCATTCCTTGACCTTGGCGAAAT |
| race Y FLAP | 32 | TTACACTCGGACCTCGTCGACATGCATTGCTATCCTTGTAGTCGCA |
| race Y FLAP | 33 | TTACACTCGGACCTCGTCGACATGCATTTCATCCAACCAGGCCTCG |
| race Y FLAP | 34 | TTACACTCGGACCTCGTCGACATGCATTCGAAGGTGTCGTCCTCGT |
| race Y FLAP | 35 | TTACACTCGGACCTCGTCGACATGCATTGATGTCCTCCAGCTGCTG |
| race Y FLAP | 36 | TTACACTCGGACCTCGTCGACATGCATTAGCGGACGAATATCCGCG |
| race Y FLAP | 37 | TTACACTCGGACCTCGTCGACATGCATTAGCCATGGATCTGCTGGT |
| race Y FLAP | 38 | TTACACTCGGACCTCGTCGACATGCATTCCTCAGGCGGAAACGCAC |
| race Y FLAP | 39 | TTACACTCGGACCTCGTCGACATGCATTACCGCGTCACCATAGTGT |
| race Y FLAP | 40 | TTACACTCGGACCTCGTCGACATGCATTTGGGTCCTGTCTCGGAGA |
| race Y FLAP | 41 | TTACACTCGGACCTCGTCGACATGCATTGCCCAATAGGTGCATGGG |
| race Y FLAP | 42 | TTACACTCGGACCTCGTCGACATGCATTCACTGCTGTGCCCACATG |
| race Y FLAP | 43 | TTACACTCGGACCTCGTCGACATGCATTCGATGTCCGCAATCTCTG |
| race Y FLAP | 44 | TTACACTCGGACCTCGTCGACATGCATTCTTCTCCGGAAAGGGGGA |
| race Y FLAP | 45 | TTACACTCGGACCTCGTCGACATGCATTAGCGCTCACATCCACCAG |
| race Y FLAP | 46 | TTACACTCGGACCTCGTCGACATGCATTTAGCCCTGCTTTTCCATC |
| race Y FLAP | 47 | TTACACTCGGACCTCGTCGACATGCATTGGAACATTTTGAGTGGCG |
| race Y FLAP | 48 | TTACACTCGGACCTCGTCGACATGCATTGAAGAAGTCGTCGCCCAT |
| sog_570 | 1 | ATTCGATGGCGTTCGATTTC |
| sog_570 | 2 | CAAGCAGACGATCAGCAGTC |
| sog_570 | 3 | AATTCGCGCAAAACTTTGCC |
| sog_570 | 4 | TACATAACTCCGAAGGGTGG |

| | | |
|---|---|---|
| sog_570 | 5 | CCACACATTCACACTTGATG |
| sog_570 | 6 | GGGCACTCGTTTTTGATATT |
| sog_570 | 7 | GAGATGGGATCATCGCATTT |
| sog_570 | 8 | TACATCCGTATCGTTTCGAT |
| sog_570 | 9 | TAGCAACGCAGCGTAATGTT |
| sog_570 | 10 | TGAGGAAATAGGAGGTGCGG |
| sog_570 | 11 | TACATGGACTTCATTTCCTC |
| sog_570 | 12 | CACATTCTGCGGATTGTAGG |
| sog_570 | 13 | TTGTGGAACAGGAAACGGGC |
| sog_570 | 14 | GCGATGAGGTGTAGAAGGAG |
| sog_570 | 15 | AATTGAATGGCACGCGGACG |
| sog_570 | 16 | GATAACACCCGCATCATCAA |
| sog_570 | 17 | TGATAGACACTGAGAGTGCC |
| sog_570 | 18 | GCAGAATGCGCTTGTAATCA |
| sog_570 | 19 | GGAGGACAACATGGAGACGA |
| sog_570 | 20 | AACTGAACAACTCCGTCTGC |
| sog_570 | 21 | CATTGAAGACCAGGGTGAGA |
| sog_570 | 22 | GCTCAATTTTCACACTCAGT |
| sog_570 | 23 | CACACGTGGAATCTCATCGA |
| sog_570 | 24 | GTATGGAAATGGGCGACGAC |
| sog_570 | 25 | ACGCGACATCAGTCGAAGAT |
| sog_570 | 26 | AGATGTGGGTACTTCTTGGA |
| sog_570 | 27 | TCTGGAAGATTTCGCAGCTG |
| sog_570 | 28 | ATCCATCGGTGTTCAAGTAG |
| sog_570 | 29 | CAAACTGATGTTGGGCCTAT |
| sog_570 | 30 | TGGTTGAAGTTGAAGCTCGG |
| sog_570 | 31 | AACTTCTCCACACTACCAAT |
| sog_570 | 32 | ATTGCACTCGTTGTCAATGG |
| sog_570 | 33 | CGTTGAATTCCTCGAGCAAT |
| sog_570 | 34 | AAGAAGCCTTCCAGATAGGA |
| sog_570 | 35 | TGGAATGGACCTCCAGATAG |
| sog_570 | 36 | AGCAGAAGCTGTTTGGAGTG |
| sog_570 | 37 | AACATTGTTGTCCGTGTAGA |
| sog_570 | 38 | CCGTTCACCAAATGGTTATC |
| sog_570 | 39 | CGATTCGTTGTAGAAGCGTC |
| sog_570 | 40 | CACATCTGACAGGAATCCTG |
| sog_570 | 41 | CATTCACCATCACGCTGAAG |
| sog_570 | 42 | GAGGATTGCGCTGAATAGTC |
| sog_570 | 43 | CAGGAATGGATGCCAACTGG |
| sog_570 | 44 | CTTCTTGTCTGGACGATAGG |
| sog_570 | 45 | TCTTGTGTCCATTGGAACTG |

| | | |
|---|---|---|
| sog_570 | 46 | CAGCACATTGGGATTGTTCG |
| sog_570 | 47 | TTCTCGTACACCTTGTTGAC |
| sog_570 | 48 | TGGGACATCAGGATCGGATG |
| sog Z FLAP | 1 | CCAGCTTCTAGCATCCATGCCCTATAAGATTGTAGGTGGTGTACATGG |
| sog Z FLAP | 2 | CCAGCTTCTAGCATCCATGCCCTATAAGTTGTGGAACAGGAAACGGGC |
| sog Z FLAP | 3 | CCAGCTTCTAGCATCCATGCCCTATAAGGCGATGAGGTGTAGAAGGAG |
| sog Z FLAP | 4 | CCAGCTTCTAGCATCCATGCCCTATAAGATAACACCCGCATCATCAAC |
| sog Z FLAP | 5 | CCAGCTTCTAGCATCCATGCCCTATAAGTGATAGACACTGAGAGTGCC |
| sog Z FLAP | 6 | CCAGCTTCTAGCATCCATGCCCTATAAGGCAGAATGCGCTTGTAATCA |
| sog Z FLAP | 7 | CCAGCTTCTAGCATCCATGCCCTATAAGAACTGAACAACTCCGTCTGC |
| sog Z FLAP | 8 | CCAGCTTCTAGCATCCATGCCCTATAAGCATTGAAGACCAGGGTGAGA |
| sog Z FLAP | 9 | CCAGCTTCTAGCATCCATGCCCTATAAGGCTCAATTTTCACACTCAGT |
| sog Z FLAP | 10 | CCAGCTTCTAGCATCCATGCCCTATAAGCACACGTGGAATCTCATCGA |
| sog Z FLAP | 11 | CCAGCTTCTAGCATCCATGCCCTATAAGACGCGACATCAGTCGAAGAT |
| sog Z FLAP | 12 | CCAGCTTCTAGCATCCATGCCCTATAAGATCCATCGGTGTTCAAGTAG |
| sog Z FLAP | 13 | CCAGCTTCTAGCATCCATGCCCTATAAGCAAACTGATGTTGGGCCTAT |
| sog Z FLAP | 14 | CCAGCTTCTAGCATCCATGCCCTATAAGTGGTTGAAGTTGAAGCTCGG |
| sog Z FLAP | 15 | CCAGCTTCTAGCATCCATGCCCTATAAGAACTTCTCCACACTACCAAT |
| sog Z FLAP | 16 | CCAGCTTCTAGCATCCATGCCCTATAAGATTGCACTCGTTGTCAATGG |
| sog Z FLAP | 17 | CCAGCTTCTAGCATCCATGCCCTATAAGCGTTGAATTCCTCGAGCAAT |
| sog Z FLAP | 18 | CCAGCTTCTAGCATCCATGCCCTATAAGAAGAAGCCTTCCAGATAGGA |
| sog Z FLAP | 19 | CCAGCTTCTAGCATCCATGCCCTATAAGTTGGAGTGCTTGGAATGGAC |
| sog Z FLAP | 20 | CCAGCTTCTAGCATCCATGCCCTATAAGCGATTCGTTGTAGAAGCGTC |
| sog Z FLAP | 21 | CCAGCTTCTAGCATCCATGCCCTATAAGCACATCTGACAGGAATCCTG |
| sog Z FLAP | 22 | CCAGCTTCTAGCATCCATGCCCTATAAGGAGGATTGCGCTGAATAGTC |
| sog Z FLAP | 23 | CCAGCTTCTAGCATCCATGCCCTATAAGCTTCTTGTCTGGACGATAGG |
| sog Z FLAP | 24 | CCAGCTTCTAGCATCCATGCCCTATAAGTCTTGTGTCCATTGGAACTG |
| lacZ 670 | 1 | CTTCTTGGGGATTCCAATAG |
| lacZ 670 | 2 | AATGGGATAGGTCACGTTGG |
| lacZ 670 | 3 | GAACAAACGGCGGATTGACC |
| lacZ 670 | 4 | TTGCACCACAGATGAAACGC |
| lacZ 670 | 5 | TTCAGACGGCAAACGACTGT |
| lacZ 670 | 6 | CGCGTAAAAATGCGCTCAGG |
| lacZ 670 | 7 | TCCTGATCTTCCAGATAACT |
| lacZ 670 | 8 | GAGACGTCACGGAAAATGCC |
| lacZ 670 | 9 | ATCGCTGATTTGTGTAGTCG |
| lacZ 670 | 10 | CACCCTGCCATAAAGAAACT |
| lacZ 670 | 11 | CTCATCGATAATTTCACCGC |
| lacZ 670 | 12 | ACGTTCAGACGTAGTGTGAC |
| lacZ 670 | 13 | TAACGCCTCGAATCAGCAAC |
| lacZ 670 | 14 | TGACCATGCAGAGGATGATG |

| | | |
|---|---|---|
| lacZ 670 | 15 | CACGGCGTTAAAGTTGTTCT |
| lacZ 670 | 16 | AGCGGATGGTTCGGATAATG |
| lacZ 670 | 17 | GGGTTTCAATATTGGCTTCA |
| lacZ 670 | 18 | GATCATCGGTCAGACGATTC |
| lacZ 670 | 19 | GATCACACTCGGGTGATTAC |
| lacZ 670 | 20 | GGATCGACAGATTTGATCCA |
| lacZ 670 | 21 | CGCGTACATCGGGCAAATAA |
| lacZ 670 | 22 | AAGCCATTTTTTGATGGACC |
| lacZ 670 | 23 | TATTCGCAAAGGATCAGCGG |
| lacZ 670 | 24 | GAAACCGCCAAGACTGTTAC |
| lacZ 670 | 25 | CCTGTAAACGGGGATACTGA |
| lacZ 670 | 26 | CATACAGAACTGGCGATCGT |
| lacZ 670 | 27 | AAACTGCTGCTGGTGTTTTG |
| lacZ 670 | 28 | CGCTATGACGGAACAGGTAT |
| lacZ 670 | 29 | GTAGTTCAGGCAGTTCAATC |
| lacZ 670 | 30 | ACCCAGCTCGATGCAAAAAT |
| lacZ 670 | 31 | CTGACTGGCGGTTAAATTGC |
| lacZ 670 | 32 | TTGTTTTTTATCGCCAATCC |
| lacZ 670 | 33 | ACTTACGCCAATGTCGTTAT |
| lacZ 670 | 34 | AATAAGGTTTTCCCCTGATG |
| lacZ 670 | 35 | CAACGGTAATCGCCATTTGA |
| lacZ 670 | 36 | GGAAGACGTACGGGGTATAC |
| lacZ 670 | 37 | GTGGGCCATAATTCAATTCG |
| lacZ 670 | 38 | TCAGTTGCTGTTGACTGTAG |
| lacZ 670 | 39 | GGAAACCGTCGATATTCAGC |
| lacZ 670 | 40 | CACCAGACCAACTGGTAATG |
| lacZ 670 | 41 | GCCCGGTTATTATTATTTTT |
| lacZ 670 | 42 | CTTACGCGAAATACGGGCAG |
| lacZ 670 | 43 | TCCTTCACAAAGATCCTCTA |
| lacZ 670 | 44 | TTGTCCAATTATGTCACACC |
| lacZ 670 | 45 | AGTTCCATAGGTTGGAATCT |
| lacZ 670 | 46 | CATTAAAGGCATTCCACCAC |
| lacZ 670 | 47 | ATGGCATTTCTTCTGAGCAA |
| lacZ 670 | 48 | CTCTTCTTTTTTGGAGGAGT |
